# Hierarchical coupling between ATP hydrolysis and Hsp90’s client binding site

**DOI:** 10.1101/2020.02.15.950725

**Authors:** Steffen Wolf, Benedikt Sohmen, Björn Hellenkamp, Johann Thurn, Gerhard Stock, Thorsten Hugel

**Affiliations:** Biomolecular Dynamics, Institute of Physics, University of Freiburg, Freiburg, Germany; Institute of Physical Chemistry, University of Freiburg, Freiburg, Germany; Engineering and Applied Sciences, Columbia University, New York, USA; Signalling research centers BIOSS and CIBSS, University of Freiburg, Freiburg, Germany

## Abstract

Several indicators for a signal propagation from a binding site to a distant functional site have been found in the Hsp90 dimer. Here we determined a time-resolved pathway from ATP hydrolysis to changes in a distant folding substrate binding site. This was possible by combining single-molecule fluorescence-based methods with extensive atomistic nonequilibrium molecular dynamics simulations. We find that hydrolysis of one ATP effects a structural asymmetry in the full Hsp90 dimer that leads to the collapse of a central folding substrate binding site. Arg380 is the major mediator in transferring structural information from the nucleotide to the substrate binding site. This allosteric process occurs via hierarchical dynamics that involve timescales from pico- to milliseconds and length scales from Ångstroms to several nanometers. We presume that similar hierarchical mechanisms are fundamental for information transfer through many other proteins.

## II. INTRODUCTION

Coupling between distant regions of a protein (allosteric communication) is an important mechanism for the regulation of protein function and signalling, as it enables alteration of protein structures at active sites by small changes in a binding pocket several nanometers away.^1–4^ In spite of its importance as elementary process of cell signalling, there is surprisingly little known about the underlying dynamical process and the timescales of allosteric communication.^5–7^ Understanding of the molecular mechanisms of allostery is crucial to, e.g., elucidate the effects of point mutations in cancer development^8^ and exploit it for the design of small molecule drugs.^9^

In this work, we investigate allostery within the structure of the yeast molecular chaperone heat shock protein 90 (Hsp90). Hsp90 is highly conserved in eucariotic cells, being involved in many intracellular signalling pathways,^10,11^ serving for example as a cyclin-dependent kinase regulator, and is therefore a target protein for cancer treatment.^12,13^ Hsp90 is a homodimer consisting of two copies from a 670 amino acid protein, with each of the chains containing three domains named N, M and C (see Fig. 1). The two monomers undergo large conformational changes, which are usually classified into N-terminal open and closed states.^14^ A nucleotide binding site is found within the N-domain (see Fig. 1a). Structural changes on the Ångstrom scale such as the hydrolysis of ATP into ADP are believed to cause significant conformational changes of the full dimer.^10,11^ Several models with atomistic resolution are available for the closed state: X-ray crystallography structures of yeast Hsp90 bound to the co-chaperone Sba-1 and the non-hydrolysable nucleotide AMPPNP (PDB ID 2CG9),^14^ the endoplasmatic reticulum Hsp90 analogue Grp94 with a folding substrate-mimicking polypeptide and AMPPNP (5ULS),^15^ the mitochondrial Hsp90 TRAP1 with AMPPNP (4IPE, 4IYN, 4J0B and 4IVG)^16^ and a cryo-EM structure of the human Hsp90 dimer bound to ATP with the co-chaperone Cdc37 and the kinase Cdk4 as folding substrate (5FWK and 5FWL).^17^ Further representative conformations of the functional cycle are found in form of the partially closed Grp94 (2O1U, 2O1V, 2O1W, and 2O1T).^18^ Structures of the open state have been proposed based on molecular dynamics (MD) simulations^19–21^ and by a combination of single molecule Förster resonance energy transfer (smFRET) and MD simulations.^22^ Despite this wealth of structural information, the molecular details on how nucleotides affect the dimer and how the nucleotide hydrolysis cycle is coupled to folding client binding and processing have remained elusive.^23^

**Fig. 1.**
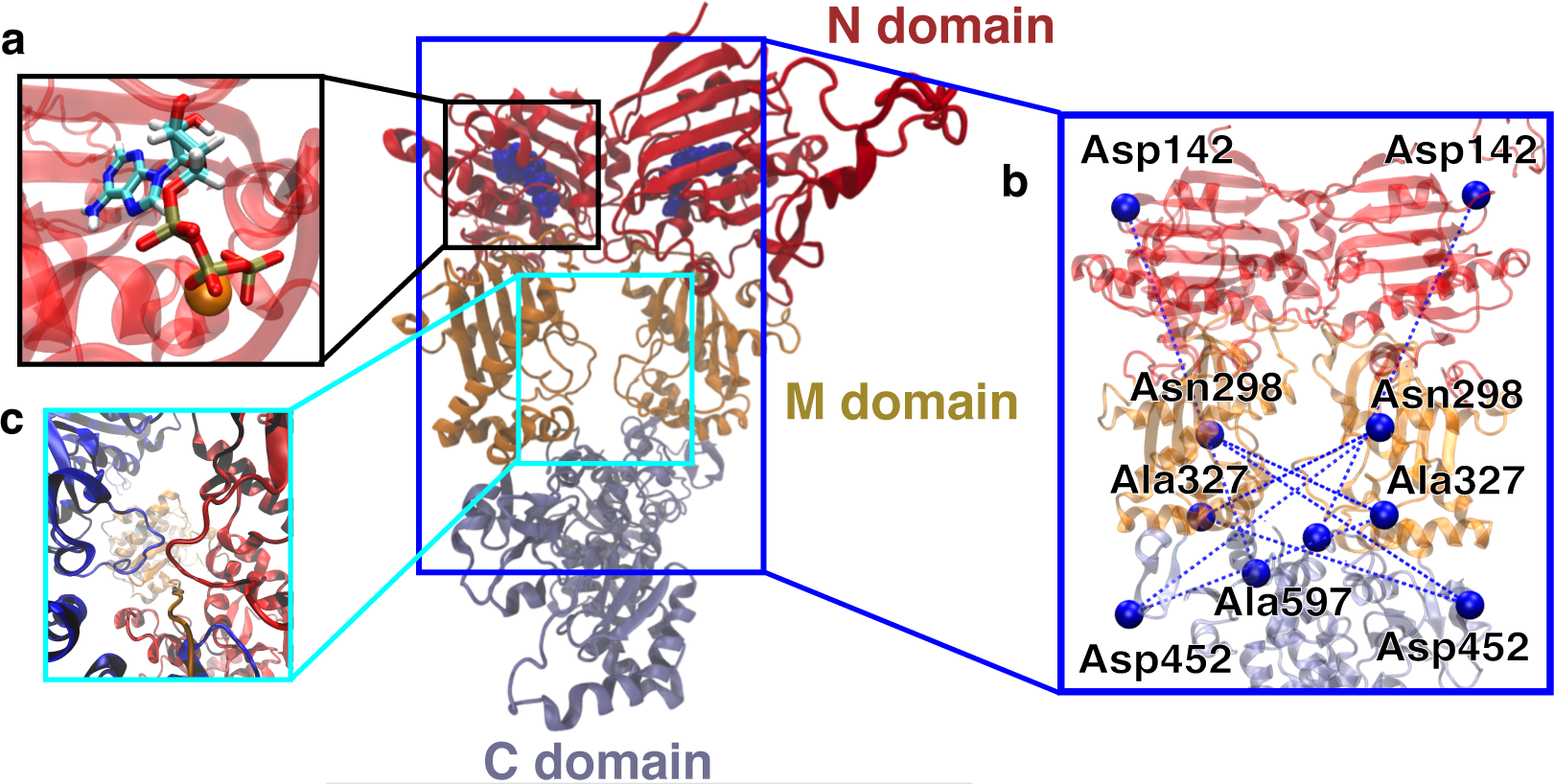
Overview of the Hsp90 dimer. The displayed structure is the initial protein model based on the X-ray crystal structure 2CG9.^14^ N-domain is shown in red, M-domain in orange, C-domain in light blue. **a** Nucleotide (in sticks) binding site in the N-domain. Magnesium ion as orange sphere. **b** FRET distance pairs. **c** Overview of the central folding substrate binding site. Protein dimer (blue and red) and Cdk4 (orange) in the active folding complex in the cryo-EM structure 5FWK.^17^

Here, we report on a joint experimental and theoretical study combining smFRET, fluorescence correlation spectroscopy (FCS) and fluorescence lifetime measurements with fully atomistic MD simulations to investigate allosteric communication in Hsp90 across several time- and length scales. Intensity-based smFRET has been used to determine conformational dynamics and distances in (bio)polymers for more than 20 years.^24,25^ More recently, single molecule lifetime and anisotropy measurements have been added to increase the range of accessible timescales and characterise physicochemical properties of fluorophores.^26,27^ We use both methods to determine distances between selected amino acid side chains in the M-domain of Hsp90 and investigate distance changes upon addition of different nucleotides. Complementary, we perform unbiased and biased all-atom MD simulations of the full dimer in a physiological NaCl solution and loadings of different nucleotides (total simulation length of 25 *µ*s) to obtain insights into molecular mechanisms which are not directly accessible with smFRET.^28–30^ We find that Arg380, which has been assumed to be involved in the functional cycle before,^31–34^ is a prominent residue for the transfer of allosteric information. Experimentally and computationally observed time- and length scales of allosteric communication agree well with each other, which allows us to propose a detailed molecular pathway from hydrolysis to a large conformational change of the M-domains.

## III. RESULTS AND DISCUSSION

### Intermolecular distance measurements

To describe the effect of nucleotides on the closed conformation of the Hsp90 dimer, we performed smFRET experiments. We attached pairs of Atto550 and Atto647N fluorophores to all combinations of the amino acid positions 298, 327 and 452 as well as the amino acids 142 and 597 (see the blue network in Fig. 1b). Three out of four of the labels are located in the middle domain of the protein dimer, i.e., between M-M and M-C domains at a distance of 3.0-5.0 nm from the nucleotide binding site. 142-597 is a complementary N-C distance pair. Each FRET label pair was measured in the apo form and additionally in the presence of 2 mM ATP, ADP, AMPPNP or ATP*γ*S (see Supplementary Methods). We note that while the protein contains ADP+P_i_ after hydrolysis, we observed no difference between ADP and ADP+P_i_ measurements (cf. Supplementary Fig. 7). In a previous study^22^ we focussed on the open ADP bound state and the closed AMPPNP state, and determined the dynamics of the Hsp90 domains on the millisecond timescale. Here we focus on the N-terminal closed states of Hsp90 under various nucleotide conditions. Choosing suitable FRET pairs, we can now distinguish two closed states of Hsp90. Using information from fluorescence correlation experiments we also found position-specific dynamics on the low-µs timescale, which can be connected to timescales observed in MD simulations.

Several of the investigated protein/nucleotide complexes clearly show three populations of preferred distances. These populations can be attributed to the existence of two closed protein structures (termed closed states A and B) in addition to the open protein conformation (see Fig. 2a,b). Even for the nucleotide conditions with a low populated closed state B, fits with three populations represent the data well (Fig. 2c). The two closed populations are consistent with several previous studies on yeast Hsp90.^37–39^ The closed states can be best distinguished in the presence of AMPPNP where they are most populated. Therefore we used the mean FRET efficiencies of the three states from the AMPPNP data to constrain fits of other nucleotides. The width of the fits was not constrained as these three global states are dynamic and might show nucleotide dependent fine structure due to small structural differences (see below). The averaged apparent distances between the dyes *R*_*(E)*_^27^ and their distributions are extracted by Photon Distribution Analysis (PDA)^35,36^ for separation of distance distributions from shot-noise (see Supplementary Methods for details and Supplementary Table 1 for all measured distances).

**Fig. 2.**
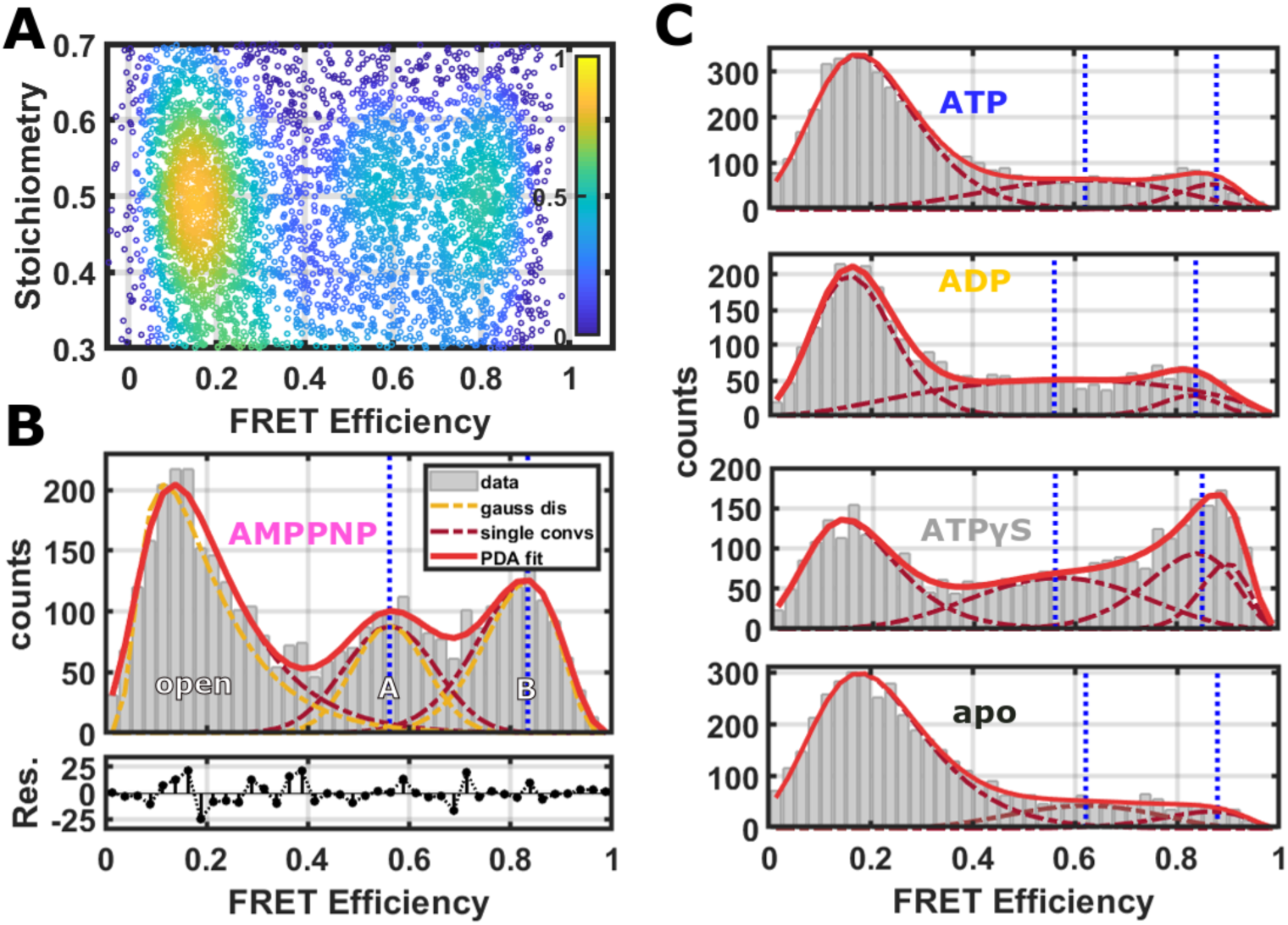
Single-molecule FRET data analysis exemplarily shown for the distance 298-452 (see Supplementary Fig. 5 for distances 298-327, 327-452 and 142-597 and Supplementary Fig. 6 for examples of the full diagrams) **a** Corrected 2D FRET efficiency vs. stoichiometry histogram in the Hsp90 dimer with AMPPNP. A kernel density estimator was used to visualise the burst density by colour. **b** FRET efficiencies between the dyes and their distributions are extracted by Photon Distribution Analysis (PDA),^35,36^ which includes a separation from shot-noise. For the 298-452 distance the PDA fit (red line) results in three states (dark red dashed lines): an open state, a closed state, termed closed state A and a more contracted closed state termed closed state B. For each population the mean distance *µ* was obtained from the shot-noise filtered distributions (orange lines) as expectation values (indicated by vertical blue dashed lines). **c** Each FRET efficiency was measured under five different nucleotide conditions. Here, PDA fits with three fixed mean distances between the dyes are shown. As the FRET efficiency (Forster radius) differs for different (swopped) dye pairs, even when the distance between the dyes is the same, blue dashed lines for the three states correspond to the same distance for the different nucleotide conditions.

To access molecular details of the population shifts observed in smFRET, we performed MD simulations of the Hsp90 dimer with different nucleotides bound (Supplementary Tab. 2). We checked if simulations resulted in a defined structural ensemble by assessing the time traces of the C_*α*_ root mean square displacement (RMSD) of the initial secondary structure elements within the full dimer (excluding the charged loop at sequence positions 208 to 280) and the radius of gyration of the full protein. As an example, Supplementary Fig. 8 displays the structural development of the global protein in a representative ATP-bound simulation. For all simulations, the initial development away from the starting structure appears to be completed after 0.5 µs.

### Nucleotide binding

As a first step towards an understanding of the structural differences imposed by the nucleotides onto the protein, we focused on structural changes found directly at the nucleotide binding sites in simulations. Figure 3a shows a representative structure of ATP bound after 1 µs simulation time. ATP is stably connected to the protein at Asp79 via the adenine moiety. The magnesium ion is *α, β, γ*-coordinated via the triphosphate, and bound to the protein via Asn37, which in turn forms a hydrogen bond with P_*α*_. The P_*γ*_ phosphate forms a stable salt bridge with Arg380. In our simulations, Arg380 is positioned directly at the tetrahedron surface of P_*γ*_ that needs to be accessed by an attacking water molecule d thus Arg380 in this position effectively shields the triphosphate from hydrolysis. This shielding could explain earlier experimental findings that ATP bound to Hsp90 in either the open or closed protein conformation exhibits a life time in the range of several seconds.^40^

**Fig. 3.**
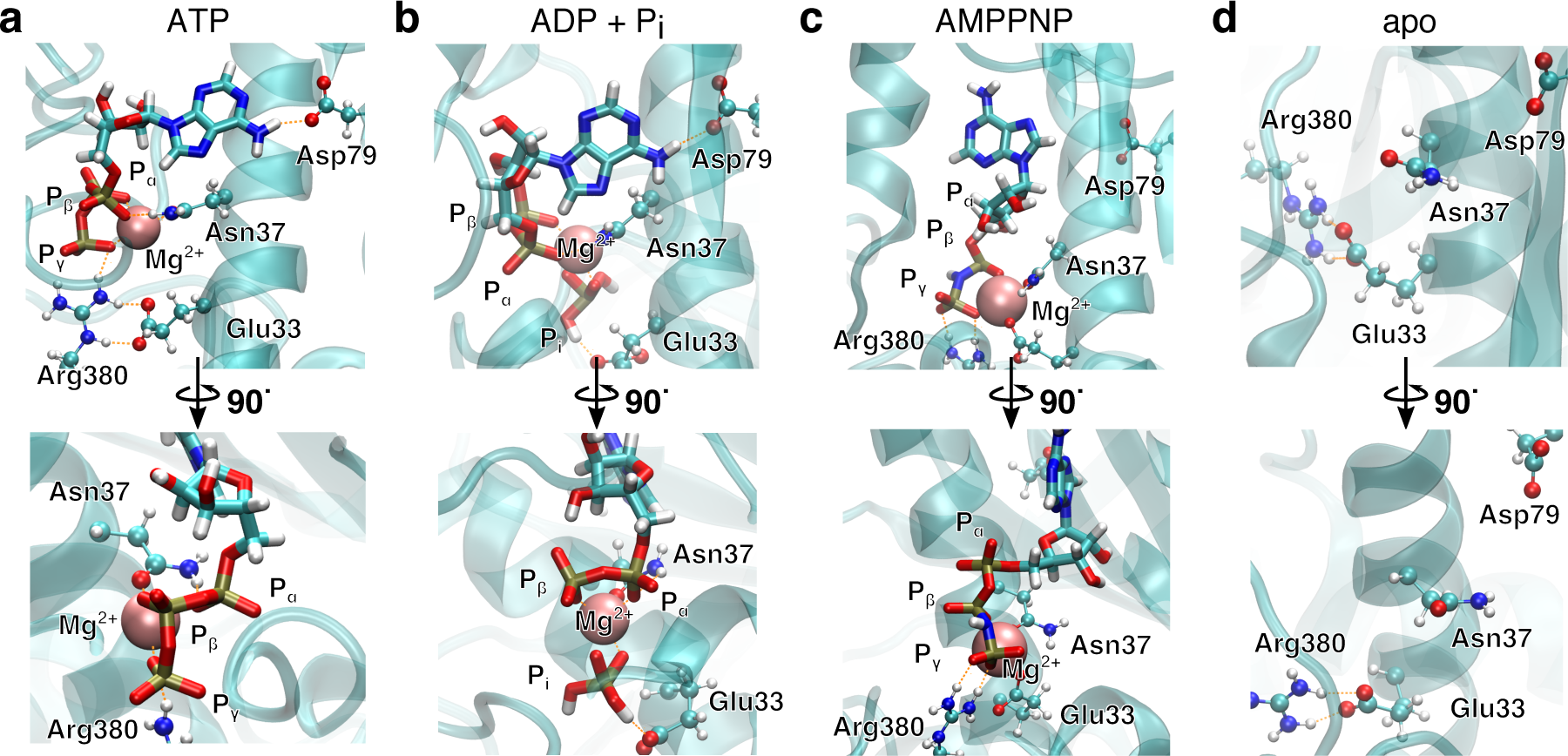
Nucleotide binding site in **a** ATP-bound, **b** ADP + Pi-bound, **c** AMPPNP-bound and **d** apo-Hsp90 simulations after 1 µs. Nucleotides displayed as sticks, amino acid side chains as balls and sticks, protein backbone as cartoon, hydrogen bonds as orange dashes. In the case of ADP + Pi, Arg380 has left the nucleotide binding site.

In comparison to this arrangement, we observe significant deviations in the simulations with ADP + P_i_ (see Fig. 3b). In the case of the doubly hydrolysed state with two ADP + P_i_ (in the following abbreviated as ADP), the binding mode of the adenine moiety and the magnesium ion is unaffected. P_*α, β*_ still coordinates the magnesium ion, but P_i_ serves as additional bidentate coordination partner. Its position interferes with the positioning of the Arg380 side chain, pushing it away from P_*α, β*_. Furthermore, the free phosphate disrupts a salt bridge of Arg380 and Glu33. This particular glutamate has been shown to be involved in the ATPase activity of Hsp90, as well.^41,42^ Arg380 in turn is highly conserved within the family of Hsp90 and its homologues^23^ and has been reported as a major player in ATP hydrolysis^20,43^ and speculated to take part in allosteric information transfer.^32^ Previously, Arg380 has indeed been considered as a potential residue to mediate the stabilisation of a specific N/M arrangement.^31^ Furthermore, it was shown that a R380A mutant completely suppresses the formation of the closed state.^33,34^ Finally, in the apo form, the binding site exhibits (at least transiently) a collapsed state, moving all mentioned protein residues closer together. Glu33 and Arg380 again are found in a salt bridge, but the position of Arg380 is significantly different from the one in the nucleotide-bound state.

Interestingly, the binding mode of AMPPNP (Fig. 3c) is significantly different from the one of ATP. The ribose moiety in one of the two monomers is flipped completely by about 180°. The binding mode of the adenine moiety of AMPPNP becomes instable in three out of five simulations (see Supplementary Tab. 2), and the P_*γ*_-Arg380 salt bridge ruptures in one simulation. Furthermore, only P_*β*_ and P_*γ*_ coordinate the magnesium ion. This effect appears to be caused by the presence of a new hydrogen bond donor in form of the NH group between P_*β*_ and P_*γ*_ that forms contacts with surrounding polar residues, as well as due to changes in charge distribution within the triphosphate resulting from the heteroatom substitution. Instead, Glu33 becomes part of the coordination sphere of the magnesium ion, disrupting the Glu33-Arg380 salt bridge.

To investigate the effect of the hydrolysis of only a single ATP molecule, we performed simulations with one ATP and one ADP + P_i_ bound, respectively (in the following abbreviated as ATP/ADP). In this case the binding mode of the adenine “anchor” residue of ADP is stable in three out of the five simulations (see Supplementary Tab. 2), while in the remaining simulations one or both anchors lose their contact with Asp79, respectively.

### Structural changes of the Hsp90 dimer

To connect smFRET measurements and MD simulations, we calculated the expected distance distributions *P* (*R* _⟨*E*⟩_) for fluorophore pairs directly from intraprotein dye-accessible volumes of the simulated structures (see Supplementary Methods). In this way, we can directly compare simulations with the measured mean distances and uncertainties.

Figure 4a shows the means and uncertainties extracted from the experimental FRET efficiency histograms (see Fig. 2, Supplementary Fig. 5 and Supplementary Methods for details). State A occurs at distances comparable to the ones found within the crystal structure 2CG9, state B represents a structure in which the protein has significantly contracted, and does not correspond to any previously structurally described state. At the start of the simulation we observe a qualitative agreement with experiment for distances attributed to the closed states A under all nucleotide conditions. In the case of ATP and AMPPNP, the simulations mainly stay in state A. However, in the presence of ATP/ADP and even more clearly in the presence of ADP, the distributions show some tendency towards closed state B. Missing convergence of MD simulations to state B indicates structural relaxation timescales longer than the maximal single trajectory length of 1 µs.

**Fig. 4.**
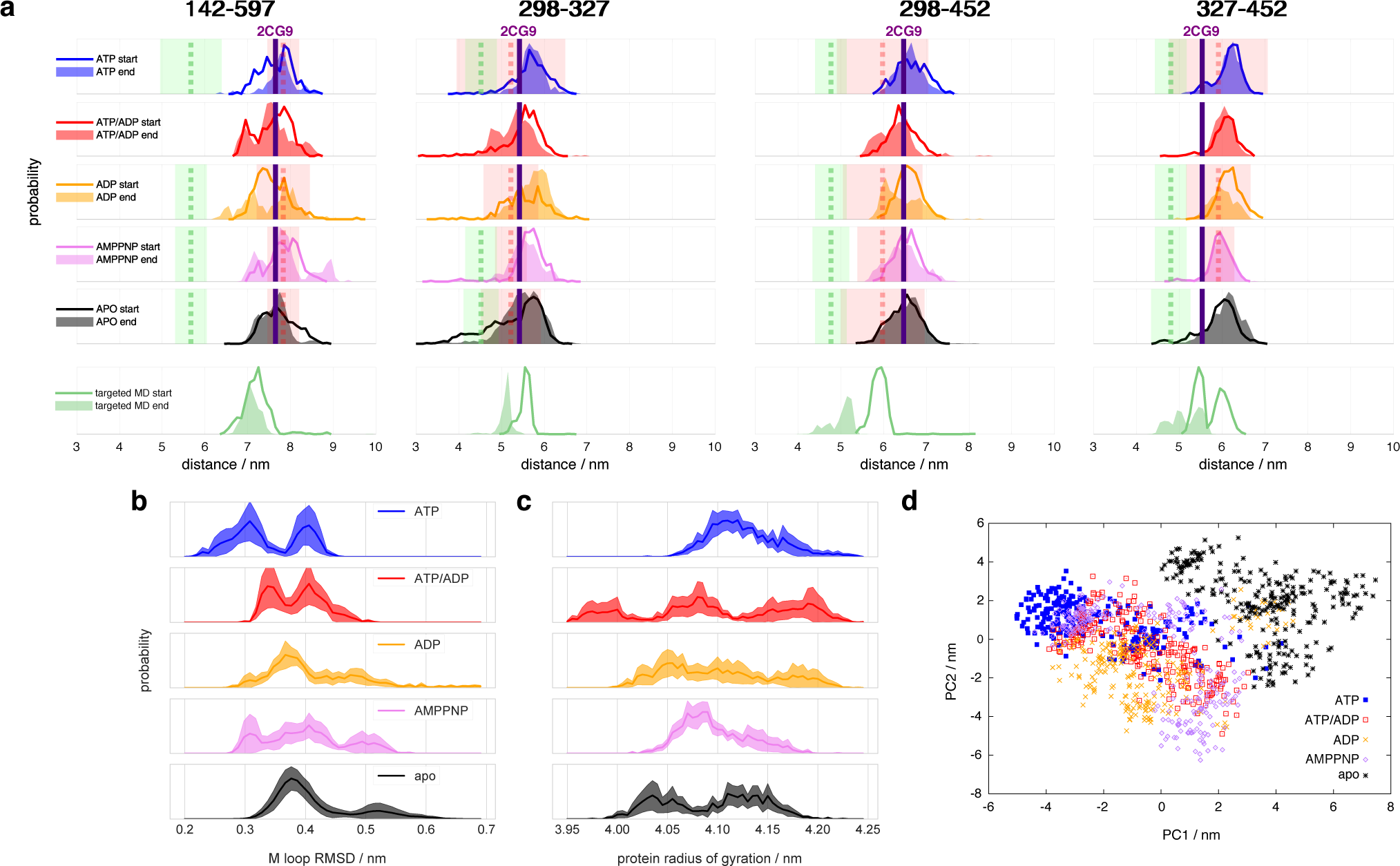
Histograms of selected Hsp90 distances. **a** Experimental and simulated apparent distances *R*_⟨*E*⟩_ of the FRET label pairs 298-327, 298-452, 327-452 and 142-597 (see Supplementary Methods for details). Line histograms display the first 0.25 µs of simulation (termed “start”), filled histograms the last 0.25 µs (termed “end”). Vertical dashed red and green lines indicate the measured mean distances (*R*_⟨*E*⟩_) of closed states A and B, respectively, and coloured transparent areas the respective uncertainties from experiment (see Supplementary Tab. 1). The vertical purple line indicate the 2CG9 structure state distance.^14^ Targeted MD represent the first and last 0.25 ns of calculations, respectively, starting from an ADP state after 1µs of unbiased MD. **b** RMSD of internal M-domain loop (sequence positions 323 to 340) during the last 0.25 µs with reference to positions in the active folding complex determined via cryo-EM (5FWK).^17^ **c** Radius of gyration of the full protein during the last 0.25 µs of simulation. **d** Principal component (PC) analysis of the last 0.5 µs of simulations of intraprotein contact distances within nucleotide binding sites and M-loops.

To accelerate structural changes leading to closed state B and to overcome the timescale limitations of unbiased simulations, we employed nonequilibrium targeted MD simulations^44^ with implicit solvent for one of the ADP simulations and pulled the *C*_*β*_ distances of amino acid pairs 298-327, 298-452 and 327-452 towards each other (see Supplementary Methods for details). The targeted MD distance distributions indeed evolve towards the experimentally measured distances (see Fig. 4a, bottom). The *R*_⟨*E*⟩_ of these three pairs agree well with their experimental counterparts. The 142-497 distance pair, which we did not manipulate, also tends towards its closed state B.

We observe that ATP generally tends to narrow distance distributions and larger overall distances, indicating that closed state A is most stable with ATP (before hydrolysis) and opens a central hole between the monomers (see below). All other states (even including AMPPNP) tend to wider distributions and/or cover shorter distances that develop towards closed state B, i.e., towards closing the hole.

Does this change in M-M distances have a functional consequence at other sites of the protein? To address this question, we compared our MD results to the Hsp90-Cdk4 cryoEM-structure (PDB ID 5FWK).^17^ Fig. 1c shows that the peptide chain of the partially unfolded client Cdk4 passes right through the core of the dimer between the two M-domains. The chain is held in place by two loops coming out of the M-domains (sequence positions 323-340).^17^ To compare the M-loop structures appearing during simulation to the corresponding cryo-EM structure, Fig. 4b displays the respective C_*α*_-RMSDs. The ATP-bound structure appears to switch between two conformations, one of which (RMSD ≈ 0.3 nm) is closest to the cryo-EM structure loop arrangement. Interestingly, AMPPNP resembles more the hydrolysed states than ATP. We further note that these M-loops are found at a distance of ca. 4.0 nm, i.e., quite far away, from the nucleotide binding site.

To assess structural differences of the full protein effected by different nucleotides, we also analyse the differences in the radius of gyration of the full dimer in Fig. 4c. The ATP-bound protein tends to a clearly defined large radius of gyration, while all other investigated states display a broader distribution and trend to smaller radii. Again, AMPPNP states resemble more the hydrolysed states. ATP binding apparently causes Hsp90 to adopt an extended closed conformation when the M-domains are considered. Such an extension in combination with a restriction of the accessed conformation space would lead to both a decreased solvent and conformational entropy. The resulting entropic penalty could be compensated by the enthalpic contributions of bound ATP, with hydrolysis leading to a return into a wider range of closed conformations with a decreased radius of gyration.

### Allosteric communication between nucleotide binding site and the protein dimer

To identify the key amino acids responsible for the structural information transfer, we performed a correlation-based principal component analysis (PCA) on the contacts^45^ made by amino acids forming the nucleotide binding site and the M-loop (see Supplementary Methods for details). The principal components display linearly independent combinations of inter-residue distances weighed according to their individual contribution to explain the correlation of distance changes in the analysed set of contacts. Hence, they can serve as “reaction coordinates” to elucidate the mechanism of structural information transfer. Figure 4d displays trajectories of the various systems projected onto the resulting first two principal components. The principal components clearly separate apo- and ATP-states. AMPPNP, ATP/ADP and ADP partially cover the ATP- and apo-states, but also areas that neither correspond to the latter two states, which is in qualitative agreement with our results given above.

Investigating the importance of individual contacts on the first two principal components (Supplementary Fig. 9), we find that besides changes within the binding pocket and the M-loop, Arg380 is highlighted to exhibit pronounced changes as found above. Arg380 is indeed prominent with respect to allosteric communication, as it is the only residue which reaches into the nucleotide binding site, but is also part of the M-domain.

On the basis of this insight, Fig. 5a shows the key structural elements of a mechanistic model of allostery. Arg380 forms a salt bridge with the P_*γ*_ of ATP, and is positioned in a loop at the N-terminal end of a helix (sequence positions 376-408, in the following called M-helix), which is part of the M-domain. The helix runs across the face of the central *β*-sheet of the M-domain, and thus can apply tension and affect the position of the whole domain. The location of the M-domain position in turn affects the orientation of the M-loop and thus regulates the formation of the folding client binding site. In other words, Arg380 serves as a “piston” to connect ATP and the N-terminal end of the M-helix via electrostatic interactions. The M-helix in turn represents a “cross-beam” that exerts force on the M-domain and effects the formation of the folding client binding site.

**Fig. 5.**
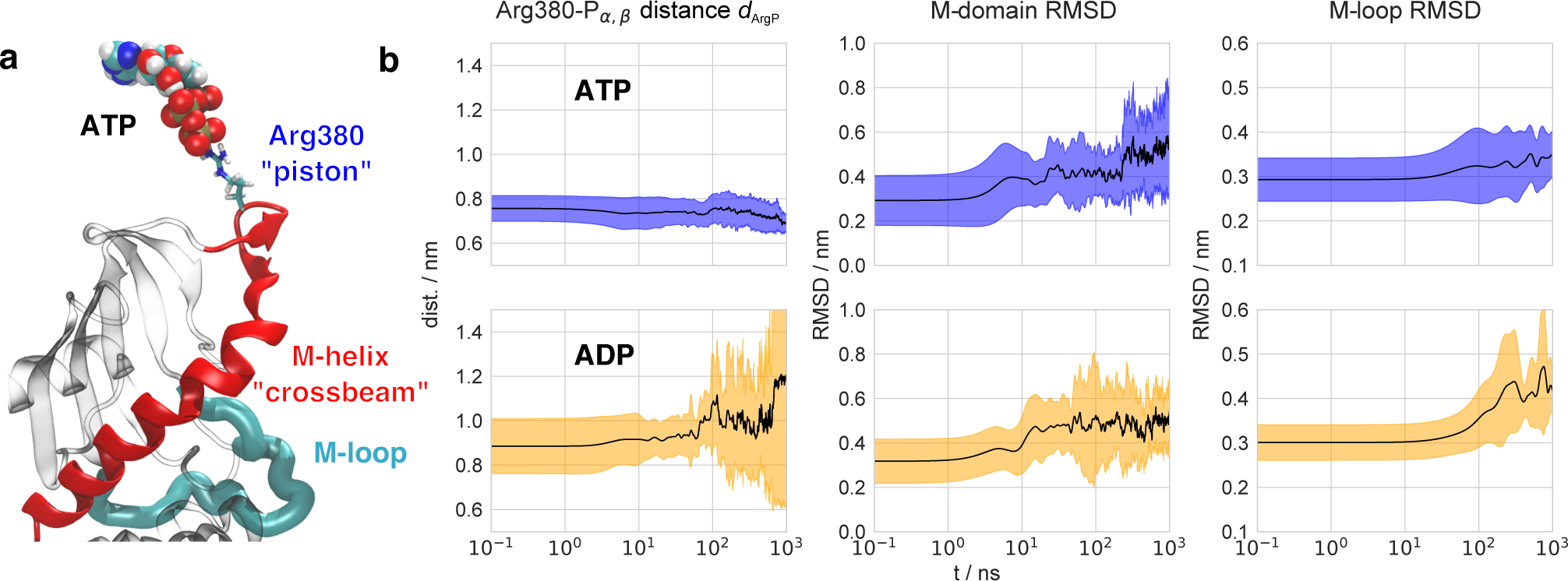
Transfer of structural changes from the nucleotide binding site to the full protein dimer. **a** Essential mechanistic coupling of nucleotide binding site and the M-domain. ATP as van der Waals spheres, Arg380 “piston” as sticks, M-domain helix “crossbeam” (residues Leu374 to Glu408) in red, M-loop in cyan, remaining M-domain in grey (other monomer not shown). **b** Time evolution of average structural changes observed in ATP and ADP state simulations, respectively. Mean values displayed in black lines, standard deviations as coloured traces. Displayed observables are: distance between the Arg380 CZ atom (center atom of guanidinium group) and the respective nucleotide P_*α*_/P_*β*_ mass center; RMSD of the C*α* atoms from secondary structure elements (helices, sheets) in the M-domain after fit of the respective N-domain C*α* atoms from secondary structure elements (helices, sheets); RMSD of the two internal M loops in respect to the conformation of the substrate bound cryo-electron microscopy structure (PDB ID 5FWK).^17^

It is instructive to consider the effect of this mechanistic coupling on the different nucleotide states (Supplementary Fig. 10a-c). In the ATP-bound state, we observe that the Arg380–P_*γ*_ salt bridge is present in both monomers. Arg380 pulls on the M-helices in both monomers, resulting in a very symmetric dimer structure. In the case of ATP/ADP, on the other hand, the free phosphate stays within the binding site and disturbs the position of one of the two Arg380 side chains. In this case, Arg380 moves away from the nucleotide, relieving the tension applied to the M-helix in the respective monomer. The Arg380 side chain in the other monomer with ATP still pulls in its M-helix. The resulting imbalance in connectivity between both monomers introduces a structural asymmetry, which has been observed to exist in other experiments.^16,46^ In the case of full hydrolysis (ADP), the retraction of Arg380 from the nucleotide binding site is present for both monomers, likely recovering structural symmetry. We emphasise that the structural symmetry of the protein seen here is not effecting a symmetry in ATP hydrolysis. ATP hydrolysis appears to be a stochastic event, leading to the ATP/ADP state as the most probable post-hydrolysis structural intermediate. Hydrolysis appears to happen independently in both monomers, and thus a single hydrolysis event is the most likely one. To support this assumption, we have compared the symmetric dimer E33A-E33A with the asymmetric dimer E33A-WT. As can be seen in Supplementary Fig. 11, the former remains closed in the presence of ATP, while the latter is mainly found in the open state.

It is interesting to note that in the proposed mechanism, the major role of ATP is the addition of electrostatic interactions to the chaperone, stabilising a client-binding competent conformation via induced structural stress. This hypothesis is in line with the observation that ATP exhibits a comparatively long life time in the binding pocket.^40^ Hydrolysis then returns the chaperone into an inactive state with a wide range of accessible conformations and, from a thermodynamic perspective, completes the overall energetic balance for the folding process. We therefore hypothesise that ATP acts as a regulatory element activating Hsp90 similar to GTP in GTPases,^47^ instead of directly performing work on the folding client.

The allosteric communication suggested here differs from the one proposed for the bacterial homolog of Hsp90, HtpG,^19^ as the involved structural elements (in particular the N-domain) differ significantly between HtpG and yeast Hsp90. Our observation that ATP hydrolysis shifts the equilibrium between the two closed states towards the more contracted state agrees well with a recent EPR study on yeast Hsp90^39^. In addition, recent simulations on human Hsp90 have implicated the loop region around Arg380 to take part in allosteric communication as well, but were too short to observe the structural changes discussed here.^48^

### Time evolution of structural changes in MD simulations

The model of allosteric communication in Hsp90 developed above suggests a sequence of information flow from the nucleotide binding site to the folding client binding site. To test this idea, we perform a time series analysis of our trajectories. Based on our findings above, the simulation starting structure based on AMPPNP-bound 2CG9 is a mixture between an ATP-like and a partially hydrolysed state. We therefore consider the ADP trajectories as equilibration simulations from a non-equilibrium starting structure after hydrolysis. To monitor intra-protein changes associated with this allosteric communication, we focus on (i) the distance *d*_ArgP_ between the Arg380 side chain guanidinium carbon atom and the P_*α, β*_ mass center, (ii) the C_*α*_-RMSD of M-domain position (using C_*α*_-atoms of the N-domain as fit reference) and (iii) the C_*α*_-RMSD of the M-loop arrangement (residues 323-340) in reference to the cryo-EM structure 5FWK. Observable *d*_ArgP_ highlights changes within the binding pocket directly after hydrolysis at a distance of ∼ 0.4 nm from the triphosphate, the M-domain RMSD protein report on conformational changes at intermediate (∼ 2.0 nm) distance, and the M-loop RMSD reflects conformational changes at the folding client binding site (∼ 4.0 nm distance).

Figure 5b displays the time evolution of means and standard deviations of these observables for ATP and ADP, averaging over 5 trajectories in each case. In the following, we discuss the apparent changes according to their timescales. A complementary visualisation of changes in ADP simulations is given in Supplementary Movie 1. The first differences appear directly at the start of simulations at *t*; ≲ 1 ns: in ATP simulations, *d*_ArgP_ is constant and exhibits only small fluctuations at all times. The simulations with ADP start with an increased *d*_ArgP_ (∼ 0.15 nm longer) and exhibit larger fluctuations, reflecting a motion of Arg380 away from the nucleotide caused by the mobile free phosphate. These differences are a result of the different nucleotide structures in the simulations, and appear directly in the initial structure minimisation. In experiment, this conformational change is expected to occur following hydrolysis on a sub-ns timescale.

The detachment of Arg380 from the nucleotide is a prerequisite for the subsequent rearrangements of the M-domain, which occur on a 10 ns timescale. In fact, at *t* ≳ 10 ns, the M-domain RMSD in the ADP system abruptly rises to ca. 0.5 nm. This change corresponds to an increased flexibility of the M-helix, followed by a rotation of the M-domains (see Supplementary Movie 1). For ATP/ADP (see Supplementary Fig. 10d), we find a similar increase in M-domain RMSD, which appears to be slowed down and completes after 100 ns, pointing to a residual stabilisation of the active folding conformation from the single bound ATP. Indeed, mutants of Hsp90 have been reported that only allow the delayed hydrolysis of a single ATP molecule, but still retain the capability to fold substrates.^49^ We mention that also in the ATP simulations an initial increase in M-domain RMSD to ∼ 0.4 nm takes place at similar times, followed by a delayed jump to a value similar to the one in the ADP simulations at ca. 300 ns. We interpret this change to come from the ATP simulations undergoing structural changes towards the folding-competent state. This involves conformational changes that are different from the one in the ADP simulations, but apparently follow a similar sequence of events.

The rotation of the M-domains is a prerequisite for the collapse of the folding client binding site, which is indicated by the M-loop RMSD and appears for *t* ≳ 100 ns. While the mean RMSD increases to ca. 0.35 nm in ATP, it reaches ca. 0.45 nm in ADP, which is in agreement with the distance-resolved RMSD profile in Fig. 4b. Lastly, *d*_ArgP_ strongly increases after *t* = 0.8 µs in ADP and exhibits large fluctuations. On longer timescales (*t* > 1 µs), these overall fluctuations may coincide with a destabilisation of the N-N interface and thus the opening of the dimer.^22^

The observed process timescales apparently follow a logarithmic scale, which is a hallmark of hierarchical dynamics, where fast processes associated with small free energy barriers regulate slow transitions:^50,51^ changes of the Arg380 position within; ≲ 1 ns cause M-domain rearrangements on a 10 ns scale, which in turn are followed by sub-µs conformational changes of the M-loops forming the folding client binding site. The missing convergence to state B-associated distances implies that the full transfer from state A to the proposed state B takes even longer. Indeed, already the relaxation of small proteins from nonequilibrium states involves timescales of tens of microseconds and beyond,^7^ and thus is outside of the scope of current unbiased MD simulation techniques.

### Timescales observed from lifetime and FRET-FCS data

Motivated by this *in silico* timescale analysis, we investigated on which timescales dynamics can be observed in our experimental data. First we tested for dynamics on the ms timescale by FRET efficiency vs. fluorescence lifetime plots.^52,53^ Fig. 6 reveals signatures of dynamics on the ms timescale between the closed states A and B. This is consistent with the analysis shown in Fig. 2, because the additional population for the dynamic intermediate state between states A and B is only little populated. These experimental findings are also consistent with the MD simulations, which do not show a full transition from closed state A to B within the timescale of the simulation.

**Fig. 6.**
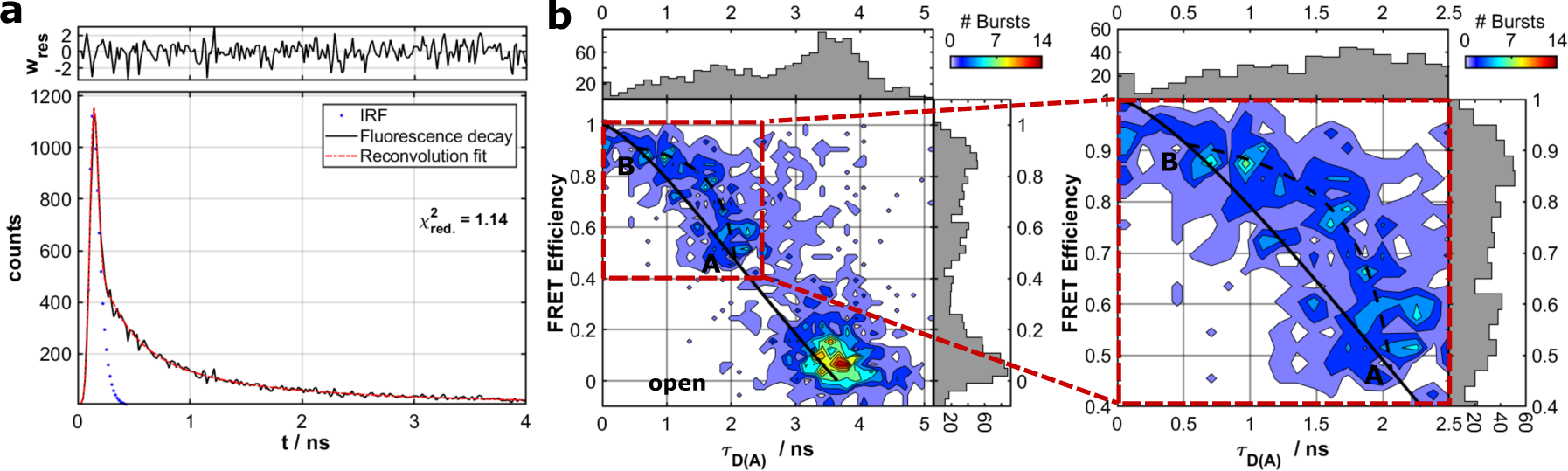
(a) Subensemble lifetime analysis of the Hsp90 closed state B for the variant 452-298 with AMPPNP. *τ*_D(A)1_=0.52 ns and *τ*_D(A)2_=2.1 ns are obtained from a reconvolution approach with a biexponential model function. Fluorescence intensity of the donor channel after donor excitation is shown as black line, the reconvolution fit as red line and the instrument response function (IRF) in blue. (b) FRET Efficiency *E* vs. donor lifetime *τ*_D(A)_ for the Hsp90 variant 452-298 with AMPPNP. A completely static sample should lay on the theoretical static FRET line (straight black line). Lifetimes obtained from the reconvolution fit were used to determine start and end point of the dynamic FRET line (dashed black line). The zoom into the region of closed state A and B clearly shows, that the data are better described by the dynamic FRET line which hints towards millisecond kinetics between closed state A and B (see Supplemental Methods for more details).

To further test if some very fast dynamics, comparable to the MD simulation timescale, can be seen, we performed FRET-FCS measurements.^54–56^ We calculate the autocorrelation and cross-correlation functions of the donor fluorescence and the FRET-induced acceptor signal (see Fig. 7). A thorough analysis (see Supplementary Methods) suggests correlation times on three timescales. The first correlation time on the order of a few milliseconds corresponds to the diffusion time of Hsp90. The two other correlation times most likely correspond to structural dynamics within the Hsp90 dimer. The estimated timescales of the found dynamics are in the range of *τ*_*L*_ ≈ 50 − 200 µs and *τ*_*K*_ ≈ 1 − 4 µs and are shown for two FRET pairs and all nucleotide conditions in Supplementary Fig. 12–15. The efficiency-lifetime analysis hints towards an additional slower rate (larger than 1ms). The fast rates are likely caused by the dynamic ensemble of asymmetric structures that are observed in the MD simulations, which finally leads to the closure of the folding substrate binding site (closed state B). The slower rates are likely due to the interconversion between closed state A and B.

**Fig. 7.**
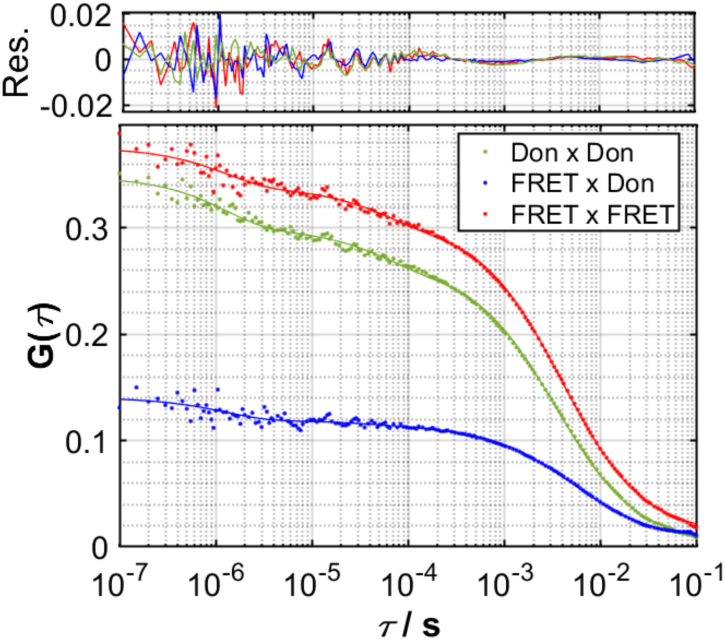
Microsecond kinetics revealed by FRET-FCS. FRET-FCS analysis for the FRET pair 298-452 in the presence of AMPPNP. Correlations of the parallel and perpendicular donor signal (green), FRET signal (red) and the cross-correlation of the parallel FRET signal with the perpendicular donor signal (blue) are shown. A global fit with two kinetic terms revealed dynamics on the timescale of about 1*µs* and 60*µs*, respectively, with a weight of around 10%, consistent with the lifetime-efficiency analysis (see Supplementary Figs. 12–15 for details and additional data).

While the presented channel of allosteric communication transfers structural information from the nucleotide to the M-domains, it only forms a small part of possible allosteric interactions within the dimer. For example, it is known that Trp300 appears to transfer structural information on folding client binding to the M-domains.^57^ Intriguingly, Trp300 is found on a loop domain that lies exactly on the opposite face of the central *β*-sheet of the M-domain to the one facing the M-helix. We speculate that client contacts with Trp300 thus may assist the M-helix in pulling the M-domains into the folding-capable conformation, and antagonize the structural compaction upon ATP hydrolysis presented here. Another recently reported allosteric switch for the regulation of Hsp90 function is found in Lys594 at the C-C domain interface.^58^

### Structural model of state B from targeted MD simulations

To obtain a model for the structural changes underlying *τ*_*L*_, we evaluate the data from our biased MD simulations (see Supplementary Methods for details). The structural consequences are displayed in Supplementary Fig. 16 and in the Supplementary Structure set: one of the monomers gains a kink at its M-C interfaces and bends over towards its partner monomer, resulting in an asymmetric contracted state, which we tentatively attribute to represent a structural model of closed state B. This asymmetric state, in which one of the two monomers significantly differs in its form from its neighbor, has so far not been described.

## IV. CONCLUSIONS

We are now in a position to formulate a model of allosteric communication involving Arg380 and the M-helix depicted as steps (1)-(6) in Fig. 8: starting with ATP (1), Arg380 forms a salt bridge with P_*γ*_, and the resulting electrostatic interaction is transmitted via the M-domain to the M-loops, keeping the folding substrate binding site open. In the following steps, sub-ns hydrolysis cleaves the connection between nucleotide and piston (2), the strain exerted by the M helix on the M-domains is lost within a nanosecond (3), and within about 10 ns the protein relaxes into an entropically favorable distribution of conformations (4) that form contracted states on a timescale of several hundred nanoseconds, where the M-domains are closer to each other and have rotated (5). Finally, on the timescale of several hundreds of µs the dimer contracts into the asymmetric state B (6). Such hierarchical timescales for the individual steps in this allosteric communication hint towards a rugged free energy landscape involving several tiers underlying this transition.^7,51^ Comparing this structural evolution with the crystal structure 2CG9, we note that 2CG9 appears as a hybrid between the ATP state (judged by the FRET distances) and the ADP state (judged by the position of the M-domains and the M-loop). This mixture of states might originate from AMPPNP and an allosteric effect of the co-chaperones bound to the N-domains in 2CG9. For the design of chemotherapeutics, novel inhibitors of Hsp90 could thus be based on creating small organic molecules that occupy the Arg380 position in the N-domain or interfere with the Glu33-Arg380 salt bridge or that pull on Arg380, similar as to the action of ATP, thereby forcing the protein into a long-lived extended state and preventing it from getting into the contracted state.

**Fig. 8.**
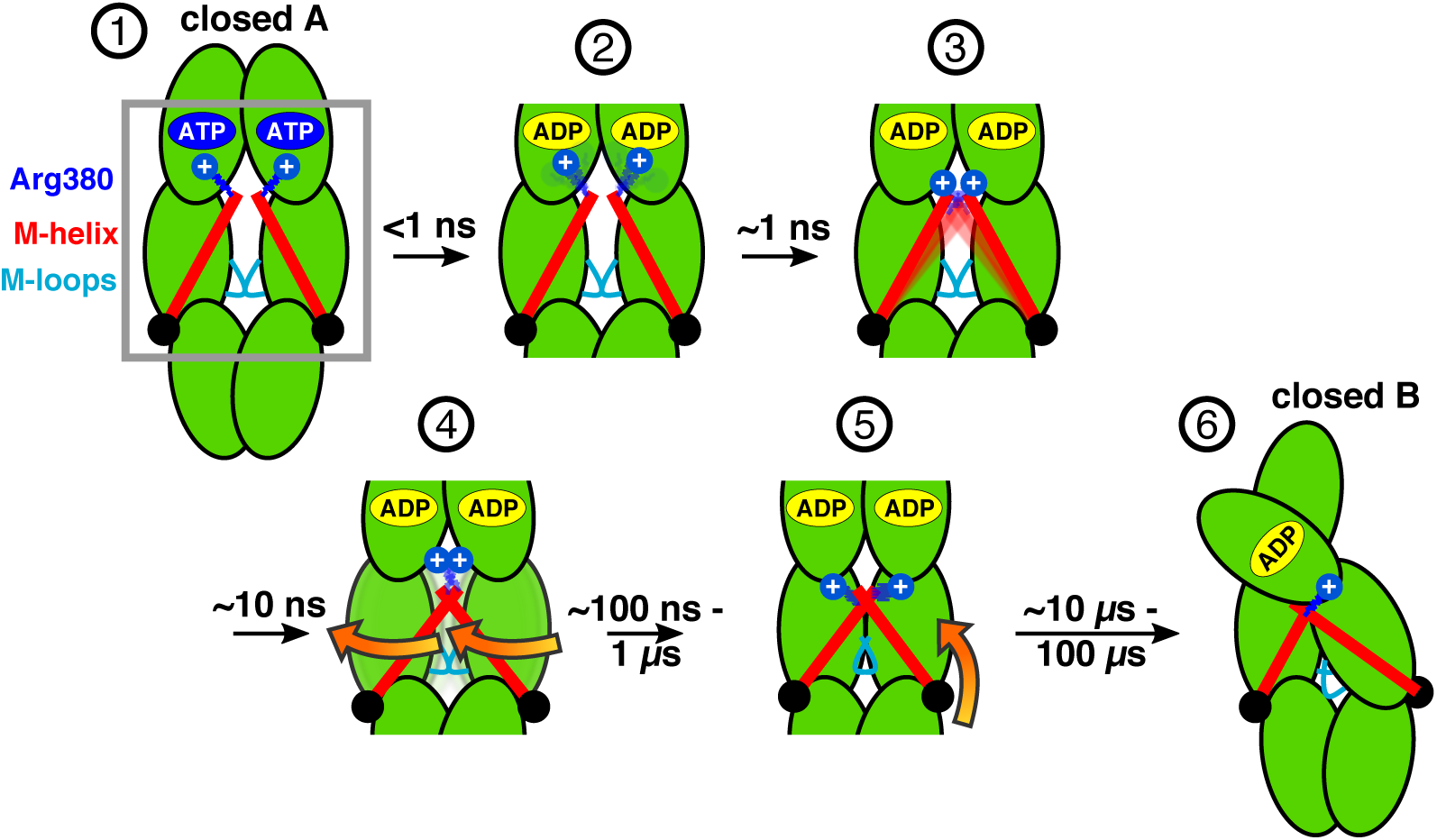
Scheme of allosteric communication including timescales. Steps 1–5 can be observed in unbiased MD simulations, steps 5 and 6 are in agreement with smFRET and FRET-FCS data and targeted MD simulations. Note that our data indicate that the hydrolysis of one ATP is already sufficient to induce the structural changes towards closed state B (see Supplementary Fig. 10). See Supplementary Fig. 16 for a structural model of the closed state B.

The presented combination of smFRET, FCS and lifetime measurements with MD simulations holds the promise to investigate time-resolved molecular mechanisms of signalling and regulation in many other biomolecular systems. Last but not least, we believe that our findings are not only key to a better understanding of the Hsp90 machinery, but will contribute to generally recognise the allosteric information transfer through proteins.

## V. METHODS

### A. smFRET measurements

Single molecule measurements were carried out on an home-build confocal microscope as depicted in Supplementary Fig. 1. FRET efficiency and apparent donor-acceptor distance *R*_⟨*E*⟩_ between the dye-pairs were determined as in^27^. FRET-FCS and lifetime data were evaluated with the PAM software.^59^ Details on Biochemistry and sample preparation, uncertainties in single-molecule experiments and the detailed analysis of the experiments are described in the Supplementary Methods and Supplementary Figs. 2 to 4.

### MD simulations and data analysis

The yeast wild type Hsp90 dimer model was created by applying MODELLER^60^ to one Hsp90 monomer (chain A) from the yeast Hsp90 crystal structure (PDB ID 2CG9).^14^ All simulations of the Hsp90 dimer water were carried out using Gromacs 2016 (Ref.^61^) using the Amber99SB*ILDN-parmbsc0-_*χ OL*3_ + AMBER99ATP/ADP force field.^62^ ATP/ADP and phosphate parameters were derived from Refs.^63^ and^64^. Missing H_2_PO_4_^−^ and AMPPNP parameters were generated based on a protocol we have used before.^65^ Details on protein modeling and the force field, the parameter generation procedure, unbiased simulations and nonequilibrium targeted molecular dynamics simulations,^44^ correlation-based contact principal component analysis (conPCA)^45^ and comparing smFRET data to MD simulation data are given in the Supplementary Methods.

## DATA AVAILABILITY

Final MD structures of the 1 *µ*s simulation runs are available with the online version of this manuscript. Additional data such as MD trajectories can be obtained from the authors upon request. The PCA software *fastpca* is available under https://www.moldyn.uni-freiburg.de/software/software.html or directly at https://moldyn.github.io/FastPCA/.

## ACKNOWLEDGMENTS

This work has been supported by the European Research Council through ERC grant agreement No. 681891, and by the German Research Foundation (DFG) under Germany’s Excellence Strategy (CIBSS EXC-2189 Project ID 390939984) and through grant No. Sto 247/10-2. The authors acknowledge support by the bwUniCluster computing initiative, the High Performance and Cloud Computing Group at the Zentrum für Datenverarbeitung of the University of Tübingen, the state of Baden-Württemberg through bwHPC and the German Research Foundation (DFG) through grant No. INST 37/935-1 FUGG, and by the Freiburg Institute for Advanced Studies (FRIAS). TH thanks Timothy Street for helpful discussions.

## AUTHOR CONTRIBUTIONS

S.W., B.H., G.S. and T.H. designed and guided research; S.W. carried out and analysed simulations; B.S. performed and analysed smFRET and FCS experiments with help from J.T. and B.H.; S.W., B.S., B.H., G.S. and T.H. wrote the manuscript.

^†^ These authors contributed equally.

## ADDITIONAL INFORMATION

### Supplementary Information

Supplementary Methods, two Supplementary Tables, sixteen Supplementary Figures, three Supplementary Notes (PDF) one Supplementary Structure set (PDB) one Supplementary Movie (MP4) are available with the online version of this manuscript.

