## Supplementary Information for "Hierarchical coupling between ATP hydrolysis and Hsp90’s client binding site"

(Dated: 1 July 2020)

### I. SUPPLEMENTARY METHODS

#### A. Biochemistry and sample preparation.

Yeast Hsp90 was recombinantly produced in *E. coli* and purified as described before.[1] Point mutations were inserted into the yeast Hsp90 gene to obtain single cysteines variants. Fluorescent labels were site-specifically introduced by cysteine-maleimide chemistry. We used ATTO550 as donor and ATTO647N as acceptor fluorophore, respectively (ATTO-TEC, Siegen, Germany). An inserted coiled-coil motif (DmKHC, *D. melanogaster*) at the C-terminus of Hsp90 prevented dimer dissociation and therefore increased the local concentration as we have shown before.[2] To obtain heterodimers which contain only one donor and one acceptor fluorophore, homodimers with the respective dye were mixed in an 1:1 ratio and incubated for 40 min at 43°C. This enabled a monomor-monomer exchange stochastically leading to 1/2 heterodimers and 1/2 homodimers. To remove aggregates, samples were centrifuged for 1h at 4°C and 16900 g. We applied size exclusion chromatography (SEC) to check that the amount of possibly remaining aggregates is sufficiently low (see Supplementary Fig. 2 for details). The biofunctionality of the new variant was tested by ATPase assays (see Supplementary Fig. 3).

#### B. smFRET measurements.

Single molecule measurements were carried out on an home-build confocal microscope as depicted in Supplementary Fig. 1. Pulsed green and red laser light (532nm, LDH-P-FA-530 and 640nm, LDH-D-C-640, respectively, PicoQuant) was polarised, overlayed and focused on the sample by an 60x water immersion objective (CFI Plan Apo VC 60XC/1.2 WI, Nikon). Excitation light was separated from the emitted light by a dichroic mirror (F53-534 Dual Line beam splitter z 532/633, AHF). The emitted light was then guided through a further dichroic mirror (F33-647 beam splitter 640 DCXR, AHF) to separate donor and acceptor fluorescence. After spectral separation pinholes with a diameter of 150$\mu$m refined the detection volume to 8fL. Finally, the two photon streams were separated by polarizing beam splitters into their parallel and perpendicular parts and recorded by single-photon detectors (two SPCM-AQR-14, PerkinElmer and two PDMseries APDs, Micro Photon Devices). Time-correlated single photon counting with picosecond resolution and data collection was performed by a

HydraHarp400 (PicoQuant) and the Symphotime 32 software (PicoQuant). To reach the single-molecule level we adjusted the protein concentration to about 50 pM. Measurements were recorded for 1800s. All experiments were carried out at 22°C in 40 mM HEPES, 150 mM KCl and 10mM $MgCl_2$ at pH=7.5. ADP and ATP were added to the protein solution directly before the measurement, in case of AMPPNP and ATP$\gamma$S, the samples were pre-incubated for 30 minutes. Nucleotides were added such that their final concentration was 2 mM.

#### C. smFRET data analysis.

Donor and acceptor photon streams were collected into 1 ms bins. Bursts were selected by a minimum threshold of 100 photons. For every burst the FRET efficiency $E$ and a stoichiometry $S$ was determined as detailed in Ref.[3]. FRET populations were determined within the region of $0.3 < S < 0.7$ of corrected $E$ vs $S$ plots. From the FRET efficiency an apparent donor-acceptor distance $R_{\langle E \rangle}$ between the two dyes is determined as[3]

$$R_{\langle E \rangle} = R_0 \left(E^{-1} - 1\right)^{\frac{1}{6}}, \tag{1}$$

where $R_0$ denotes the Förster radius. We use the efficiency-averaged apparent distance $R_{\langle E \rangle}$, because we assume that during a burst the complete accessible volume of the respective dye is sampled homogeneously, i.e., $E$ is already an average efficiency. $R_{\langle E \rangle}$ can be calculated by the FPS software[4] as is visualized in Supplementary Fig. 4. Note that $R_{\langle E \rangle}$ is also used as a variable for the average from many efficiency averaged bursts, which is only equivalent to $R_{\langle E \rangle}$ in Equation 1 if protein dynamics is negligible.

For every FRET label pair, the Förster radius $R_0$ was calculated from the donor quantum efficiency $Q_{\mathrm{D}}$, the spectral overlap $J$, and the relative dipole orientation factor $\kappa^2$

$$R_0^6 \propto \kappa^2 \, Q_{\mathrm{D}} \, J. \tag{2}$$

Here the dipole orientation factor is assumed to be $\langle \kappa^2 \rangle = 2/3$ for isotropic coupling. The position-specific donor quantum yield $Q_{\mathrm{D}}$ is given in terms of in dependence of the measured lifetime $\tau_\mathrm{D}$ as well as the parameters $Q_\mathrm{D0}$ and lifetime $\tau_\mathrm{D0}$ specified by the manufacturer[5] as

$$Q_\mathrm{D} = Q_\mathrm{D0}\frac{\tau_\mathrm{D}}{\tau_\mathrm{D0}}. \quad (3)$$

Based on the efficiency distributions (see Fig. 2 and Supplementary Figs. 5and 6) and the respective $R_{\langle E\rangle}$ distribution, we determined the expectation value $\mu$ of the distance between the two dyes and the apparent distance fluctuation $\sigma$ via a Photon Distribution Analysis (PDA).[6,7] To that end, we fitted a shot-noise-broadened sum of Gaussian distance distributions to our $E$ histogram, and performed an axis transformation from $E$-space to $R_{\langle E\rangle}$-space via Equation (1). Specifically, we transform only the bin edges. The Gaussian distance fraction of each histogram bin is obtained by integrating from the left to the right edge and extracting the mean distance. Additionally, the $E$-specific shot-noise contribution is calculated for each bin, which can be described by a beta-function. We derive its parameters from the number of photons per burst, direct excitation and leakage similar to Ref.[7]. The final convolution is gained by simply summing all bin-wise shot-noise beta-functions weighted with the value of the Gaussians for each bin. This essentially assumes a distance delta-function (no protein dynamics) for each bin that undergoes a broadening by shot-noise, which simplifies the convolution greatly. The amplitudes, mean values $\mu$ and width $\sigma_\mathrm{Gauss}$ of each Gaussian are then optimised by minimising the squared residuals between data and fit.

#### D. Uncertainties in single-molecule experiments.

The widths of the shot-noise-broadened Gaussian fits is caused by shot-noise and structural fluctuations of the protein itself. While $\sigma_\mathrm{Gauss}$ is therefore not directly related to an uncertainty of the mean value $\mu$, in our case it is a good indication for the uncertainty of the fit. In addition, in Fig. 4 we indicate a minimum uncertainty of 0.36 nm based on findings from structural remodelling.[5]

#### E. Lifetime analysis.

To investigate dynamics of the Hsp90 closed state on the millisecond timescale we performed FRET efficiency $E$ vs donor-lifetime-in-the-presence-of-acceptor $\tau_\mathrm{D(A)}$ analysis using the PAM software[8] according to Refs.[4,9,10]. Herein, kinetics can be identified by comparing the experimental burst distribution to theoretical lines: the 'static FRET line'for a completely static sample and the 'dynamic FRET line'. To obtain the endpoints of the dynamic FRET line we applied sub-ensemble lifetime analysis. Using the filters $0.3 < S < 0.7$ and $0.65 < E < 1.0$ we obtained all bursts and microtimes belonging to closed state B. The microtimes from the parallel and perpendicular donor channels after green excitation were histogrammed. The instrument response function (IRF) was obtained from a pure water measurement. Correct channel alignment was achieved by correcting for IRF shifts due to count rate dependent timing. Different detection efficiencies for parallel and perpendicular detection were considered by the G-factor (see the end of the paragraph for a summary of correction parameters). A reconvolution fit with a bi-exponential model function was applied to the fluorescence decay and provided the dynamic FRET line endpoints. Note, that we did not use constraints for the lifetimes which could explain the slight deviations between the FRET efficiency of closed state B here (0.9) and the $ES$ plot in Fig. 2a. $E$ vs. $\tau_{\mathrm{D(A)}}$ analysis was performed for the stoichiometry-filtered burst subset of the Hsp90 closed state region. The bursts are smeared along the dynamic FRET line which is evidence for millisecond dynamics of Hsp90 closed state A and B. We emphasise that the dynamic line is not the result of a fit, but an analytic solution.[9–11] Comparing the measured data to these limiting cases, the zoom in main text Fig. 6B shows that the data clearly fits better to the dynamic line. Please note that the dynamic intermediate state between states A and B is only very little populated and therefore not defined as a state in the PDA analysis of Fig. 2.

Summary of correction parameters for the lifetime fit of Hsp90 closed state B:

| **Start** | **Length** | **IRF Length** | **IRF Shift** | **Scat Shift** | **Perp Shift** | **G-factor** |
|---|---|---|---|---|---|---|
| 8 | 322 | 71 | 5.2 | 4.9 | -1.7 | 1.2 |

A reconvolution approach with a biexponential model function was applied to the fluorescence decay of the donor channel after donor excitation. Note, that for consistence we use the same correction parameters as they are declared in the PAM software[8]. 'Start'denotes start channel of the data with respect to the PIE channel start, 'Length'is the data range, 'IRF length'determines the channels which are used for IRF reconvolution, 'IRF Shift'the

IRF shift with respect to the sample fluorescence, 'Scat Shift'the shift of the scatter pattern with respect to the sample fluorescence, 'Perp Shift'accounts for differences of the IRF shift in the parallel and perpendicular channel and the G-factor considers differences of parallel and perpendicular detection efficiencies.

##### F. FRET-FCS analysis

FRET-FCS measurements were carried out in SiPEG surface-passivated chambers at 22°C. Protein concentrations were in the range of 500 pM to 1.5 nM and the buffer used was 40 mM Hepes, 150 mM KCl and 10 mM $MgCl_2$ at pH=7.5. Measurement times were 600s and measurements were repeated at least once. According to Barth et al.[12] we calculated the correlations DonxDon, FRETxFRET and DonxFRET using the PAM software[8] and Matlab R2017a. We tested several fit models, also including triplet terms (see below of an overview of the tested models). In a first approach we fixed the triplet fraction and triplet relaxation time to values that were separately determined to reduce the number of free parameters. We measured free Atto550 and free Atto647N dyes in solution and applied fit model A which contains a simple 3D diffusion term and a triplet term. However, as triplet characteristics are sensitive towards dye environments and further experimental paramters (e.g. laser excitation power), it is essentially preferable to derive them from the experiments directly. Therefore, we additionally calculated acceptor acceptor correlations (par x perp) for each data set. Obtained triplet times $\tau_T$ were in the range of 1-10$\mu s$ with fractions around 5% which is consistent with previous studies[13]. This was also expected because we used sufficiently low laser powers which were within the linear range of a laser power vs. count rate plot (not shown here).

As the triplet fraction is very low, we proceeded similar to Barth et al.[12] and we fit the experimental data without a triplet term but with a kinetic term instead (model B). We obtained a fast kinetic rate which is on the same time scale as the triplet rate but with a fraction of more than 10%. We concluded, that an additional process on this time scale must be present which is not related to the triplet state of the dyes used. Finally, best fits were obtained in model C, which includes two kinetic terms and no triplet term. Thereby we obtained a second slower kinetic rate on the hundred microsecond time-scale. Note, that the fact, that we can fit DonxDon, FRETxFRET and DonxFRET with model C whereas

AccxAcc is well described by model A, is a negative control, because for AccxAcc one would not expect conformational dynamics.

Based on the idea of separating conformational dynamics from diffusive dynamics[11] we fit DonxDon, FRETxFRET and DonxFRET with globally linked relaxation times $\tau_K$ and $\tau_L$. We obtained diffusion times $\tau_D$ in the range of 2-4 ms, depending on the analysed FRET pair and the nucleotide condition. As aggregates impede correct interpretation of correlation data we applied the software-integrated algorithm for filtering[8]. To stay consistent we applied the same aggregate filter parameters for all data sets (threshold 40, time window 10, add window 3).

**Tested fit models:**

Model A

$$G(\tau) = \frac{\gamma}{N}(1 + \frac{T}{1-T}e^{-\frac{\tau}{\tau_T}})(1 + \frac{\tau}{\tau_D})^{-1}(1 + \frac{\tau}{\rho^2\tau_D})^{-0.5} + \text{const.}$$

Model B

$$G(\tau) = \frac{\gamma}{N}(1 + Ke^{-\frac{\tau}{\tau_K}})(1 + \frac{\tau}{\tau_D})^{-1}(1 + \frac{\tau}{\rho^2\tau_D})^{-0.5} + \text{const.}$$

Model C

$$G(\tau) = \frac{\gamma}{N}(1 + Ke^{-\frac{\tau}{\tau_K}} + Le^{-\frac{\tau}{\tau_L}})(1 + \frac{\tau}{\tau_D})^{-1}(1 + \frac{\tau}{\rho^2\tau_D})^{-0.5} + \text{const.}$$

Assuming a gaussian shape of the confocal volume we set the geomatric factor $\gamma$ to $1/\sqrt{8}$. $N$ is the average number of particles in the confocal volume. $T$ and $\tau_T$ are triplet fraction and triplet relaxation time, respectively. Diffusion is characterized by the diffusion time $\tau_D$. $\tau_K$, $\tau_L$ with fractions $K$ and $L$ describe two relaxation times with corresponding fractions, respectively. The setup-related factor $\rho$ describes the ratio between the axial and lateral diameter of the confocal volume and was determined beforehand. It was measured before each experiment by scanning of matrix-immobilized polymer beads which are smaller than the diffraction limit. Values for green and red varied from 3.4 to 4.1. Furthermore, a constant was included to account for dynamics occuring at timescales exceeding the experimental observation time.

#### G. Anisotropy criterion

Sufficient rotational freedom of protein-coupled FRET dyes is a prerequisite for reliable distance measurements and can be measured by time-resolved anisotropy experiments[5]. Nucleotide- and subpopulation-specific time-resolved anisotropies were determined for each FRET pair. We did so by identifying all photons of a population from the 2D $ES$ plot, histogramming their microtimes and calculating the anisotropies $r(t)$:

$$r_{\mathrm{DD}}(t) = \frac{I_{DD}^{\parallel} - g_G I_{DD}^{\perp}}{I_{DD}^{\parallel} + 2g_G I_{DD}^{\perp}} \tag{4a}$$

$$r_{\mathrm{AA}}(t) = \frac{I^{\parallel}_{AA} - g_G I^{\perp}_{AA}}{I^{\parallel}_{AA} + 2 g_R I^{\perp}_{AA}} \tag{4b}$$

Here, the subscripts DD and AA denote photons from the green channel after green excitation and photons from the red channel after red excitation, respectively. Polarisation of the photons is indicated by the superscripts $\parallel$ and $\perp$. $I$ is the number of photons in the respective microtime bin and $g$ a detection correction parameter for green ($g_G$) and red ($g_R$) detection. The combined residual anisotropies were calculated as a geometric mean of the donor and acceptor residual anisotropies[5]:

$$r_c = \sqrt{r_{\mathrm{DD}}(t \to \infty)}\sqrt{r_{\mathrm{AA}}(t \to \infty)} \tag{5}$$

We determined the combined residual anisotropies based on the FRET-subpopulations ($r_{\mathrm{c}}^{\mathrm{a}}$), as well as based on the donor- and acceptor-only subpopulations ($r_{\mathrm{c}}^{\mathrm{b}}$). We decided to show both values because $r_{\mathrm{c}}^{\mathrm{a}}$ suffered from weak statistics. Although $r_{\mathrm{c}}^{\mathrm{b}}$ can be biased towards lower values due to remaining free dyes in solution, we believe that this is an important information, especially as the amount of free dye is very low. All values are shown in Supplementary Table 1.

#### H. Protein modeling.

The yeast wild type Hsp90 dimer model was created by applying MODELLER[14] to one Hsp90 monomer (chain A) from the yeast Hsp90 crystal structure (PDB ID 2CG9)[15] to add missing loops and revert point mutations. The dimer was then reconstituted in vmd[16] by copying the full length monomer and aligning both monomers with the protein backbone of 2CG9. ATP was introduced into the binding site according to the coordinates in 2CG9. We need to point out that this crystal structure does contain ATP coordinates, but was actually crystallized using AMPPNP. For modeling structures with bound ADP and phosphate, we manually introduced a geometry change of the ATP $\gamma$-phosphate as it should appear in a $S_N2$ nucleophilic attack by a water molecule. The resulting ADP + Pi complex thus represents a structure immediately after the hydrolysis reaction and before any relaxation of the protein.

#### I. MD simulations and data analysis.

All simulations of the Hsp90 dimer water were carried out using Gromacs 2016 (Ref. 17) using the Amber99SB*ILDN-parmbsc0-$\chi_{OL3}$ + AMBER99ATP/ADP force field,[18] which is an extension of AMBER99SB*ILDN[19–21] and contains improved parameters of ATP/ADP,[22,23] glycosidic torsions[24] and magnesium.[25] We modelled $P_i$ as $H_2PO_4^-$, as would be expected from the addition of a water molecule to $P_\gamma$. Missing $H_2PO_4^-$ and AMPPNP parameters were generated with antechamber[26] and acpype[27] using GAFF atomic parameters[28] and AM1/BCC charges[29] based on a protocol we have used before.[30] Minimum angles involving the N–H group between $P_\beta$ and $P_\gamma$ were derived from an AMPPNP structure minimised at the B3LYP/6-31G* level using Gaussian09.[31] The quality of AMPPNP simulation parameters was checked by comparison of a AMPPNP structure minimised in vacuo with the quantum mechanically minimised structure. $H_2PO_4^-$ and AMPPNP parameters can be found in the Supplementary Notes.

The simulation system consisted of a dodecahedral box of 17.5 nm side length filled with ca. 120,000 TIP3P water molecules.[32] Sodium and chloride ions were added to result in a charge neutral box with a 154 mM ion solution. We used a 2 fs time step and constrained hydrogen bonds by the LINCS algorithm.[33] Electrostatics were described by the particle mesh Ewald (PME) method.[34] Cutoffs were set to 1 nm for van der Waals interactions and a minimum of 1 nm for PME real space. Simulations were carried out in a NPT ensemble with the temperature set to 310 K and the pressure to 1 bar. Temperature control was achieved by the Bussi velocity rescaling thermostat[35] (coupling time constant of 0.8 ps), and pressure control via the Parrinello-Rahman barostat[36] (isotropic pressure coupling, coupling time constant of 0.5 ps, compressibility of $4.5 \text{x} 10^{-5}$ bar$^{-1}$).

Simulation boxes were first minimised using the conjugate gradient method with position restraints on protein, nucleotide and phosphate heavy atoms. By starting with different initial velocity distributions, five statistically independent simulation replicas were calculated for each nucleotide load investigated. After a first 100 ps equilibration simulation with position restraints, each unbiased equilibrium production simulation was run for a total trajectory length of 1 µs. Simulations with modelled hydrolysis were subjected to a second equilibration run of 100 ps trajectory length with a step size of 0.2 fs and removed restraints to allow the binding site to adjust to the presence of the free phosphate molecule.

Interprotein distances were assessed using a correlation-based contact principal component analysis (conPCA)[37,38] on the last 0.5 µs of each simulation. We took into account all minimal interresidue distances $\mathbf{d} = \{d_{ij}\}$ within and between the N-domain and M-loops that lie within a 0.45 nm cutoff in the final structures after 1 µs of all 25 simulations. conPCA builds a correlation matrix

$$\sigma_{ij} = \langle d_i - \langle d_i \rangle \rangle \langle d_j - \langle d_j \rangle \rangle / \sigma_i \sigma_j, \tag{6}$$

with distance variances $\sigma_i$ and $\sigma_j$ which after diagonalisation yields $n$ eigenvectors $\mathbf{e^{(n)}}$ that are aligned with the maximal correlation within the data set, and $n$ eigenvalues $v^{(n)}$ that determine the contribution of eigenvector $n$ to the overall correlation. The principal components $PC_n$ are then obtained via the projection

$$PC_n = \mathbf{e^{(n)}} \cdot \mathbf{d}. \tag{7}$$

Visualization of molecular data was performed with vmd.[16]

For nonequilibrium targeted molecular dynamics simulations,[39,40] we employed the PULL code from Gromacs. Prior to these simulations, we removed bulk solvent and ions from the final structure of one of the ADP+$P_i$ simulations after 1 µs simulation time, mimicking the influence of water by setting the relative permittivity $\epsilon_r = 78$ in the simulation, while lowering the overall friction of the system. For a simulation length length of 1 ns, we pulled the $C_\beta$ atoms of the distance pairs 298-327, 298-452 and 327-452 with a constant constraint velocity of 1 m/s, 2 m/s and 1 m/s, respectively, which roughly corresponds to the changes in $R_{\langle E\rangle}$ observed between closed states A and B (see Fig. 4a, main text). We then use the apparent distances of $R_{\langle E\rangle}$ as read-out parameter, as they are derived from fluorophore accessible volume, and not the $C_\beta$ distance. Additionally, we did not manipulate 142-597, but use it as a control parameter.

#### J. Comparing smFRET data to MD simulation data.

Usually, FRET experiments and MD simulations are compared by converting dye distances into $C_\beta$-atom distances, using geometric arguments concerning linker lengths and flexibility. Since the latter usually involves ill-defined assumptions, we instead compared measured and calculated $R_{\langle E\rangle}$. That is, we directly calculate the expected $R_{\langle E\rangle}$ for each simulations snapshot using the FPS software,[4] which analyses the volume a dye can access within the linker length around a given $C_\beta$-atom.[3,41] This yields the $R_{\langle E\rangle}$ distributions shown in Fig. 4a, which can be directly compared to the mean experimental distance $\mu$. The approach assumes isotropic averaging of dipoles during FRET, which can be verified via the low combined anisotropy of the FRET dyes (see Supplementary Tab. 1 and Ref.[3]). Even in case of partial anisotropic averaging, this approach is preferential over simple $C_\beta$ estimation, as the volumes accessible to the fluorophores significantly depend on the structures appearing during the MD simulation.

#### II. SUPPLEMENTARY TABLES

##### A. smFRET distance measurements

Supplementary Table 1. Results from smFRET and fluorescence lifetime analysis. For each FRET pair and nucleotide condition Förster radii $R_0$, fractions of the closed states A and B, mean fluorophore distances $\mu$, uncertainties $\sigma$ and combined residual anisotropies determined from FRET populations $r_C^a$ and donor- and acceptor-only populations $r_C^a$ are summarised. Note, that for one distance in principle two smFRET experiments can be examined which is due to swappping of the donor- and acceptor-dye position. Swapping of donor- and acceptor-dyes can have a small effect on the Förster radius because of its environment-sensitivity. As seen from the table, we performed these measurements for some distances as a check for self-consistency. For details on the determination of the combined residual anisotropies, please refer to the SI Methods part C. For details on the error determination, please refer to the Methods section in the main text.

| FRET pair | Nucleotide | $\mathbf{R_0}$ [nm] | frac$_A$ | $\mu_\mathbf{A}$ [nm] | $\sigma_\mathbf{A}$ [nm] | frac$_B$ | $\mu_\mathbf{B}$ [nm] | $\sigma_\mathbf{B}$ [nm] | $r_\mathbf{c}^a$ | $r_\mathbf{c}^b$ |
|---|---|---|---|---|---|---|---|---|---|---|
| **142-597** | apo | 5.40; 5.41 | 0.15 | 7.83 | 1.5E-3 | 0.09 | 5.67 | 0.02 | 0.20 | 0.20 |
| **142-597** | ATP | 5.40; 5.41 | 0.15 | 7.83 | 0.09 | 0.15 | 5.67 | 0.72 | 0.21 | 0.17 |
| **142-597** | ADP | 5.40 | 0.15 | 7.83 | 0.62 | 0.22 | 5.67 | 0.18 | 0.20 | 0.15 |
| **142-597** | AMPPNP | 5.40; 5.41 | 0.15 | 7.83 | 0.08 | 0.32 | 5.67 | 0.33 | 0.20 | 0.17 |
| **298-327** | apo | 6.37 | 0.23 | 5.22 | 0.70 | 0.03 | 4.52 | 0.41 | 0.24 | 0.17 |
| **298-327** | ATP | 6.37 | 0.23 | 5.22 | 1.27 | 0.03 | 4.52 | 0.01 | 0.24 | 0.17 |
| **298-327** | ADP | 6.37 | 0.18 | 5.22 | 0.64 | 0 | 4.52 | 2.61 | 0.21 | 0.15 |
| **298-327** | AMPPNP | 6.37 | 0.18 | 5.22 | 0.29 | 0.29 | 4.52 | 0.20 | 0.22 | 0.13 |
| **298-327** | ATP$\gamma$S | 6.37 | 0.17 | 5.22 | 0.29 | 0.15 | 4.52 | 0.01 | 0.21 | 0.12 |
| **298-452** | apo | 6.31 | 0.23 | 5.98 | 0.98 | 0.06 | 4.77 | 0.33 | 0.21 | 0.12 |
| **298-452** | ATP | 6.31; 6.41 | 0.23 | 5.98 | 1.07 | 0.07 | 4.77 | 0.31 | 0.25 | 0.15 |
| **298-452** | ADP | 6.31; 6.41 | 0.24 | 5.98 | 0.93 | 0.04 | 4.77 | 0.04 | 0.25 | 0.13 |
| **298-452** | AMPPNP | 6.31; 6.41 | 0.26 | 5.98 | 0.59 | 0.13 | 4.77 | 0.30 | 0.22 | 0.12 |
| **298-452** | ATP$\gamma$S | 6.41 | 0.33 | 5.98 | 0.71 | 0.33 | 4.77 | 0.43 | 0.23 | 0.14 |
| **327-452** | apo | 6.32 | 0 | 5.91 | 1.14 | 0.06 | 4.80 | 0.23 | 0.19 | 0.15 |
| **327-452** | ATP | 6.32; 6.47 | 0.07 | 5.91 | 1.15 | 0.08 | 4.80 | 0.13 | 0.24 | 0.15 |
| **327-452** | ADP | 6.32 | 0.15 | 5.91 | 0.75 | 0.14 | 4.80 | 0.16 | 0.20 | 0.09 |
| **327-452** | AMPPNP | 6.32; 6.47 | 0.15 | 5.91 | 0.27 | 0.32 | 4.80 | 0.19 | 0.21 | 0.14 |
| **327-452** | ATP$\gamma$S | 6.47 | 0.08 | 5.91 | 0.84 | 0.15 | 4.80 | 0.37 | 0.20 | 0.11 |

#### B. Simulation data

| protein | simulation No. | RMSD converged? | convergence time / ns | RMSD convergence | adenine bound? | Arg380 bound? |
|---|---|---|---|---|---|---|
| apo | 0 | yes | 100 | 4-5.5 | - | - |
| | 1 | yes | | | - | - |
| | 2 | yes | | | - | - |
| | 3 | yes | | | - | - |
| | 4 | yes | | | - | - |
| 2ATP | 0 | yes | 350 | 4,5 | ok | ok |
| | 1 | yes | 300 | 3,35 | ok | ok |
| | 2 | yes | 50 | 4.5 | ok | ok |
| | 3 | yes | 50 | 3.5 | B slightly moved | ok |
| | 4 | yes | 350 | 3.7 | ok | ok |
| ATP + ADP + $P_i$ | 0 | yes | 100 | 4.7 | jump (both) | ok |
| | 1 | yes | 200 | 4.6 | ok | ok |
| | 2 | yes | 300 | 4.3 | ok | ok |
| | 3 | yes | 250 | 5-5.5 | ok | no |
| | 4 | yes | 50 | 3.7 | jump/turn | no |
| 2ADP + $P_i$ | 0 | yes | 50 | 3-4.5 | ok | ok |
| | 1 | yes | 75 | 4 | ok | both unbind |
| | 2 | yes | 200 | 4 | ok | both unbind |
| | 3 | yes | 50 | 4-4.5 | ok | ok |
| | 4 | yes | 100 | 5-5.5 | ok | ok |
| 2MPPNP | 0 | yes | 250 | 4.5 | B jump | ok |
| | 1 | yes | 100 | 3.5 | ok | both unbind |
| | 2 | yes | 250 | 3.25 | ok | ok |
| | 3 | yes | 300 | 5.0 | B jump | ok |
| | 4 | yes | 300 | 4.0-5.0 | B jump | ok |

Supplementary Table 2. Statistics on simulation runs. Each simulation was carried out for 1 $\mu$s, except for 2ADP + $P_i$ / 1, which produced unreadable files after 0.784 $\mu$s due to a writing error.

#### III. SUPPLEMENTARY FIGURES

##### A. smFRET setup

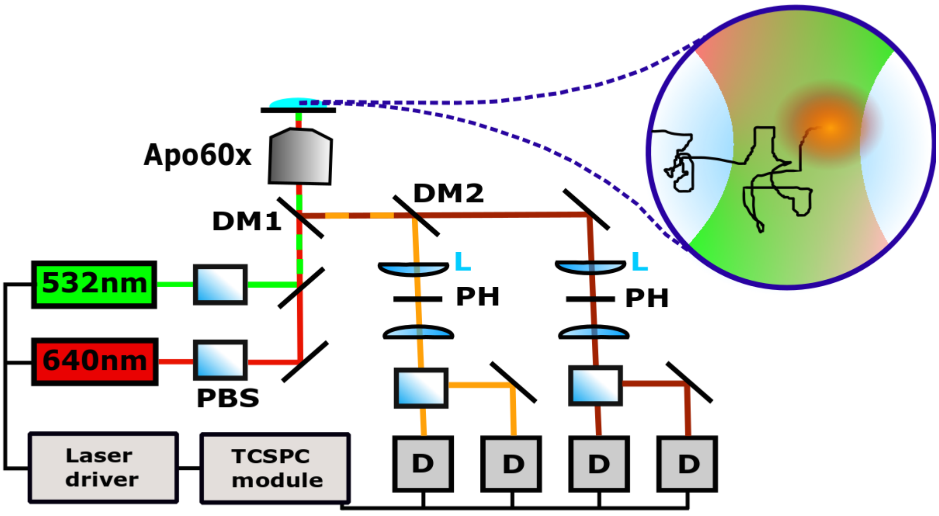


Supplementary Figure 1. Scheme of the confocal smFRET setup. Green (532 nm) and red (640 nm) laser light was polarized by polarizing beam splitters (PBS) and focussed on the sample by a water immersion objective (Apo60x). Single molecule fluorescence was separated from excitation light by a dichroic mirror (DM1), separated spectrally into donor and acceptor fluorescence (DM2) and separated with respect to polarization. Parallel and perpendicular donor and acceptor fluorescence was detected by four single photon detectors (D). Time-correlated single photon counting enabled highly time-resolved data acquisition (TCSPC module).

#### B. Sample purity controls

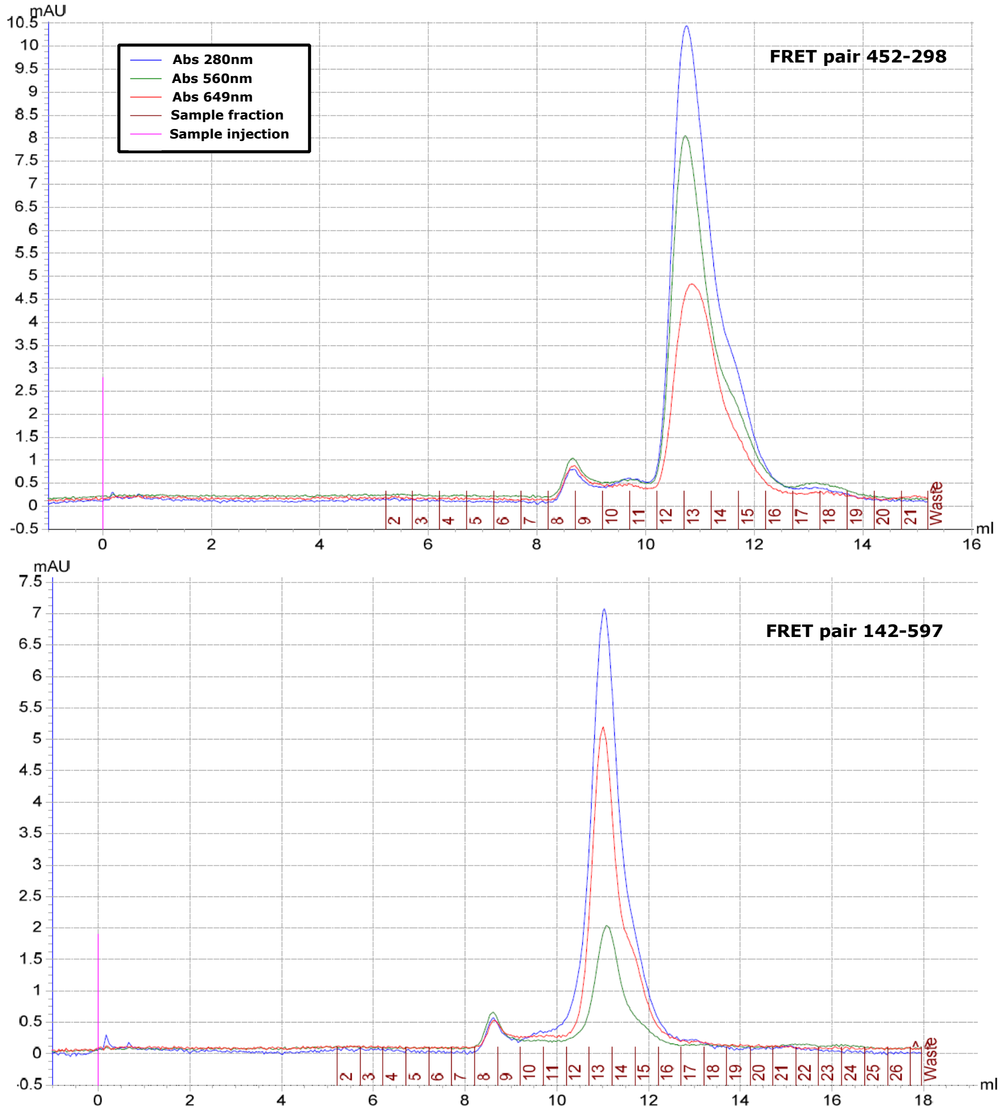


Supplementary Figure 2. SEC chromatograms for the FRET pairs 452-298 and 142-597 recorded on a Superdex 200 Increase 10/300 GL. In both cases, two main fractions are obtained with peaks at 8.3 and 13.0 ml, respectively. The first one is located within the exclusion volume of the column and therefore probably related to aggregates. We identify the second one as labelled Hsp90 dimers. The SEC profiles show, that amount of aggregates is very low with respect to the one of labelled Hsp90 dimers. Note, that in experiment, burst threshold and a stoichiometry filter further assure that aggregates are not involved in the data analysis.

#### C. Sample functionality controls

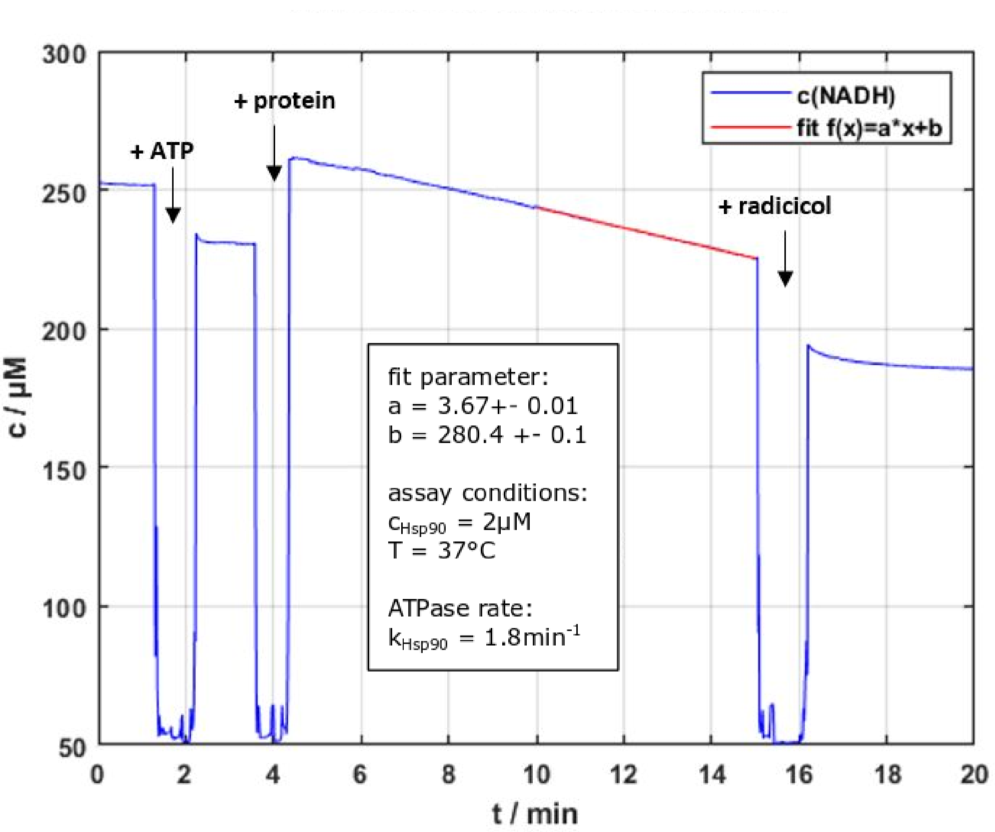


Supplementary Figure 3. ATPase activity test for the FRET pair 142-597 (all other FRET pairs have been tested and published before[5]). Hsp90 heterodimers were obtained by heating the respective homodimers, D142C and A597C, at 1:1 ratio for 43min at 43°C which promotes exchange of the monomers. Subsequent spin down (2h, 4°C, 16.9g) was performed to remove potential aggregates. The ATPase assay was performed at 37°C according to previous ATPase tests[42,43]. Absorbance at 340nm was monitored on a Lambda35 UV-VIS spectrometer (Perkin Elmer). 2 mM ATP was added to 0.2mM NADH, 10 u/ml lactate dehydrogenase, 6 u/ml pyruvate kinase and phosphoenolpyruvate solved in 40 mM HEPES, 150 mM KCl and 10 mM $MgCl_2$. The Hsp90 heterodimer 142-597 was added after the signal was stable. To determine the ATPase background, the reaction was stopped by radiciol (R2146-1MG, Sigma Aldrich) which specifically inhibits the ATPase activity of Hsp90. The ATP-turnover rate was determined from the slope of a linear fit to the decay of the signal after protein addition. We took the average from three tests and obtained $k_{Hsp90}$=(2.1±0.2) min$^{-1}$ which agrees well with previously determined values[5].

#### D. FRET label linker length correction

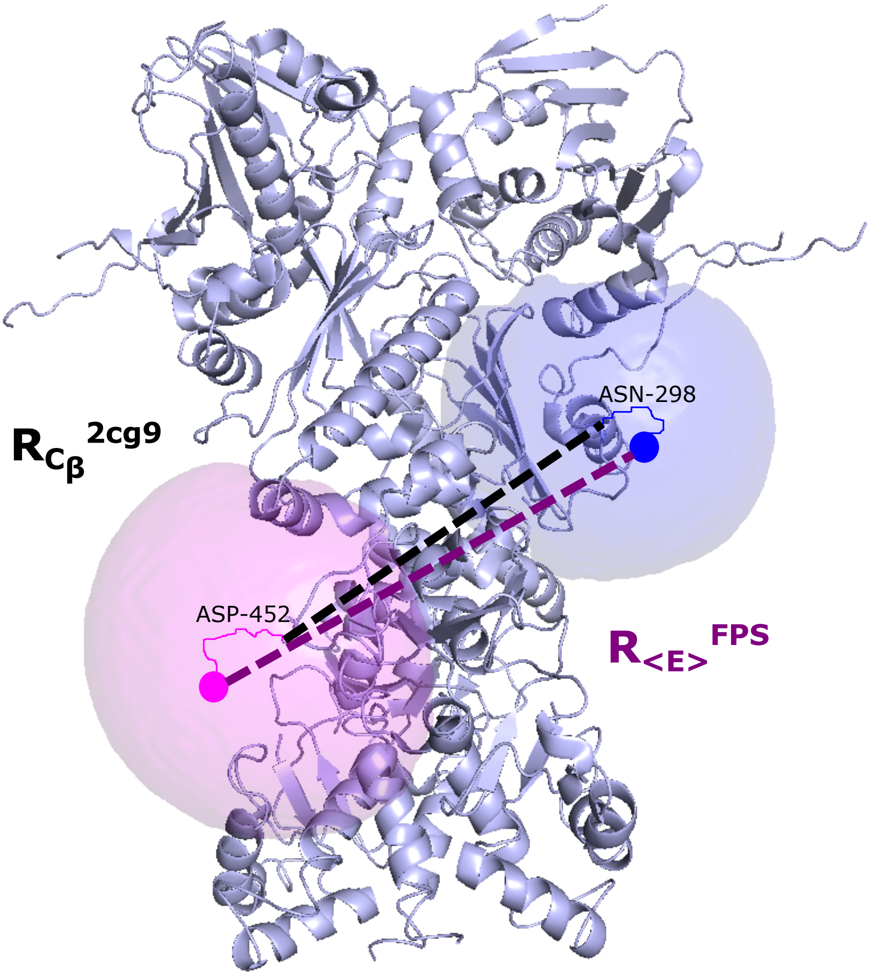


Supplementary Figure 4. Estimation of the dye linker effect. Distances obtained by smFRET and by MD simulation differ by the FRET linker which is attached to the $C_\beta$ atom of the respective Hsp90 amino acid. Here, the $C_\beta$-$C_\beta$ distance between position 452 and 298 is shown exemplarily for the 2CG9 structure (black dashed line). In order to account for the difference, for each structural MD snapshot the expected $R_{\langle E\rangle}^{\mathrm{FPS}}$ is calculated by the FPS Software.[4] Here, this is shown at the 2CG9 structure (purple line). Parameters used to calculate the accessible volumes of the dyes (blue and magenta spheres) were taken from Ref.[3].

#### E. smFRET data analysis

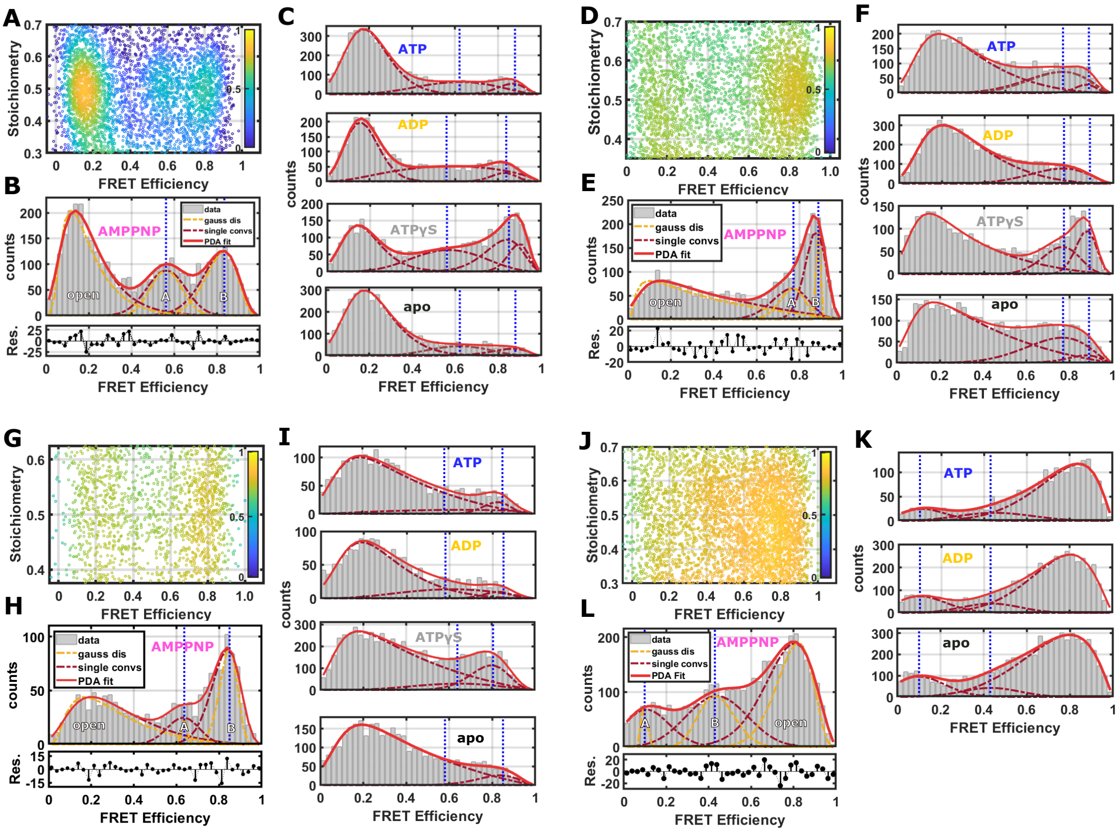


Supplementary Figure 5. Summary of single-molecule FRET analysis for all four Hsp90 FRET pairs with different nucleotides. **a-c** distance 452-298, **d-f** distance 327-298, **g-h** distance 327-452 and **j-l** distance 142-597. **a, d, g, j** Corrected 2D FRET efficiency vs. stoichiometry histogram shown with AMPPNP for 452-298, 327-298, 327-452 and 142-597, respectively. Correction of FRET efficiencies and stoichiometries was performed as described in detail in Ref.[3]. For each pair three different FRET populations were identified. A kernel density estimator was used to visualise the burst density by colour. **b, e, h, l** Distances between the dyes and their distributions are extracted by Photon Distribution Analysis (PDA)[6,7]. Under AMPPNP conditions, the three Hsp90 states emerged the most clearly. Free three-state PDA fits are shown as red lines. Superposed single states are indicated by the dashed dark-red lines and shot-noise filtered states by the dashed orange lines. Distances of each population were extracted from the expectation values of each FRET state. These are indicated for the closed state A and B by vertical dashed blue lines. **c,f,i,k** PDA analysis shows the nucleotide dependence of the FRET pairs 452-298, 327-298, 327-452 and 142-597, respectively. Investigated nucleotide conditions are ATP, ADP, ATP$\gamma$S and apo. Fits were performed with FRET efficiencies fixed to the AMPPNP distances in order to investigate the nucleotide-specific state-population. Please note that FRET efficiencies are shown here. As the FRET efficiency (Förster radius) differs for different (swopped) dye pairs, even when the distance between the dyes is the same, blue dashed lines within one distance pair correspond to the same distance.

#### F. Example full E-S plots

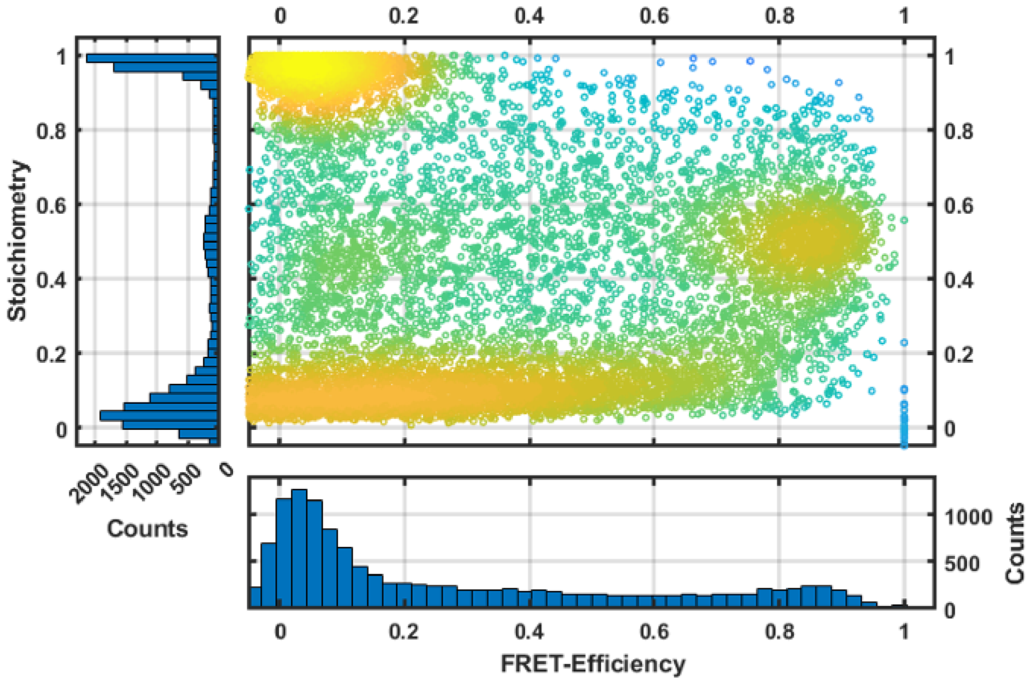


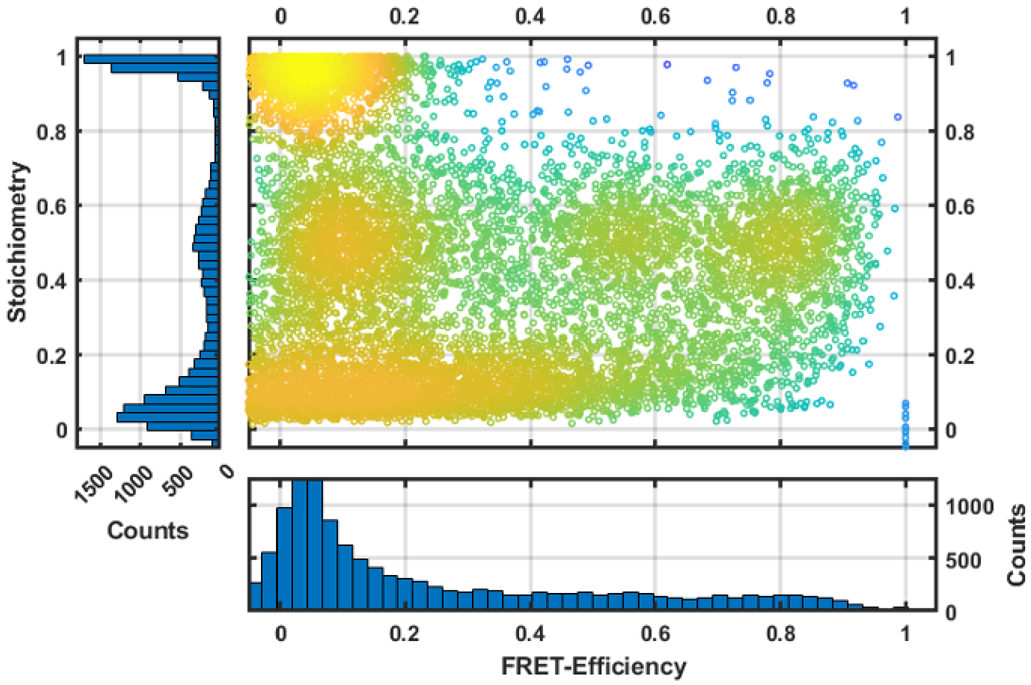


Supplementary Figure 6. full E-S plot for the variant 327-298 (top) and as a comparison for the variant 452-298 (bottom). To indicate the burst rates we added histograms on the side of the E-S plot.

#### G. ADP vs. ADP+Pi

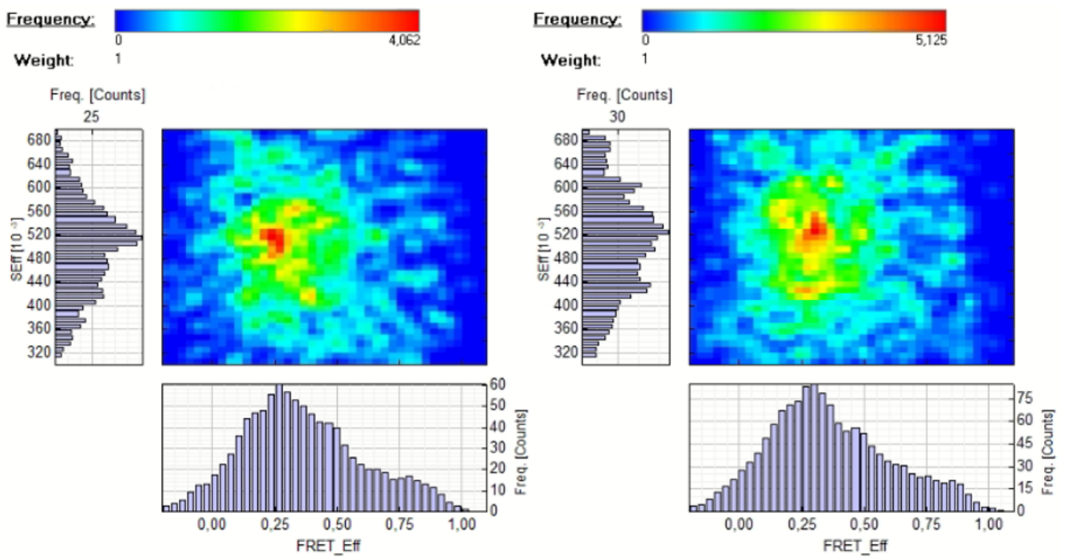


Supplementary Figure 7. ADP vs. ADP + Pi loaded Hsp90. Left: 1 mM ADP. Right: 1 mM ADP + 20 mM phosphate. Both conditions lead to near-identical FRET efficiency histograms.

#### H. MD simulation equilibration

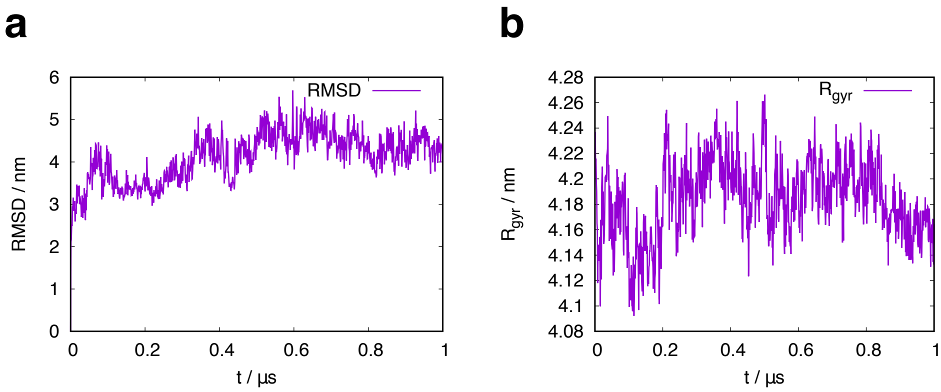


Supplementary Figure 8. Time development of **a** secondary structure $C_\alpha$ root mean square displacement and **b** global radius of gyration in a representative trajectory with ATP-bound protein.

#### I. Principal component analysis

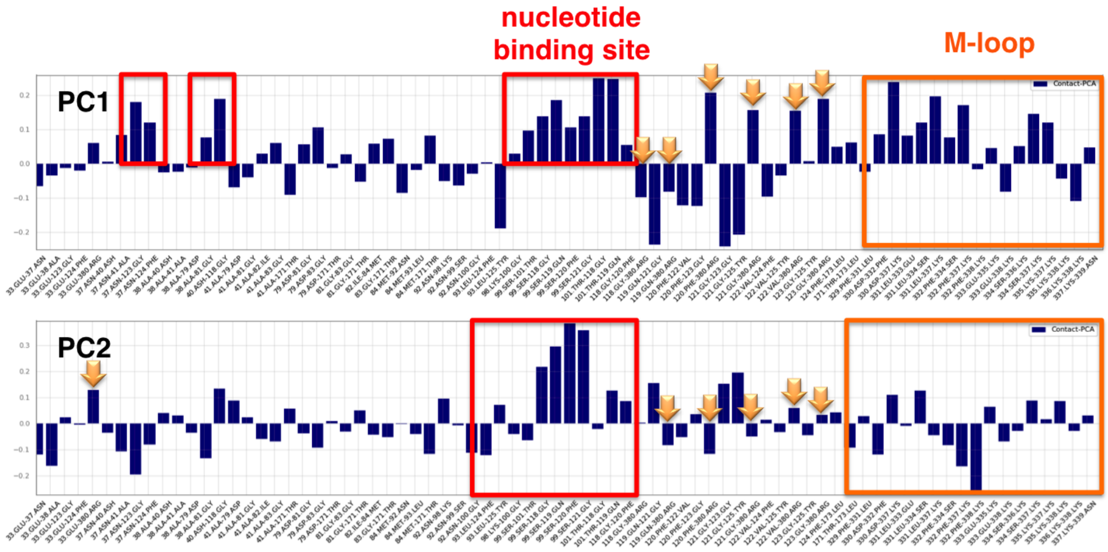


Supplementary Figure 9. Content of individual contacts within principal components 1 and 2. Nucleotide binding site contacts in red, M loop contacts in orange. Contacts of Arg380 highlighted with yellow arrows.

#### J. Effect of hydrolysis

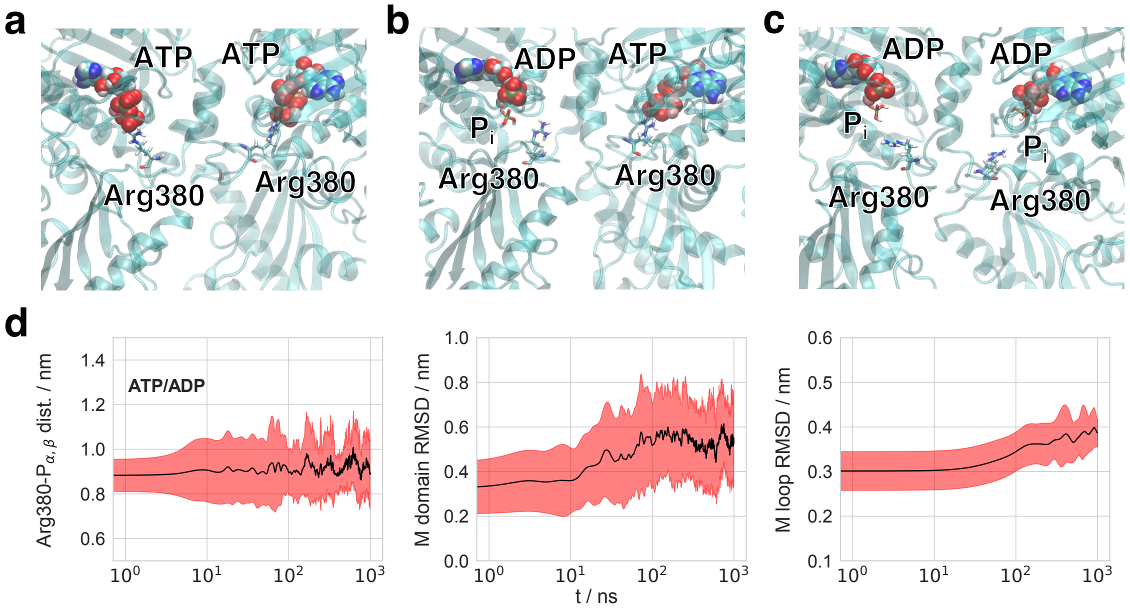


Supplementary Figure 10. **a-c** Changes of the binding site and Arg380 between ATP-bound, single hydrolysis and doubly hydrolyzed states. ATP, ADP and magnesium ion as van der Waals spheres, $P_i$ and Arg380 as sticks. Displayed are the respective structures at the end of one selected simulation for all three states (simulations "1" in Supplementary Tab. 2). **d** Time-resolved average structural changes in ATP and ADP state simulations, respectively. Mean values displayed in black lines, standard deviations as coloured traces. Displayed observables are: distance between the Arg380 CZ atom (center atom of guanidyl group) and the respective nucleotide $P_\alpha$ / $P_\beta$ mass center; RMSD of the C$\alpha$ atoms from secondary structure elements (helices, sheets) in the M-domain after fit of the respective N-domain C$\alpha$ atoms from secondary structure elements (helices, sheets) - measures the shift in position of the M-domain in respect to the N-domain; RMSD of the two internal M loops in respect to the conformation of the substrate bound cryo-electron microscopy structure (PDB ID 5FWK).[44]

#### K. Asymmetric hydrolysis

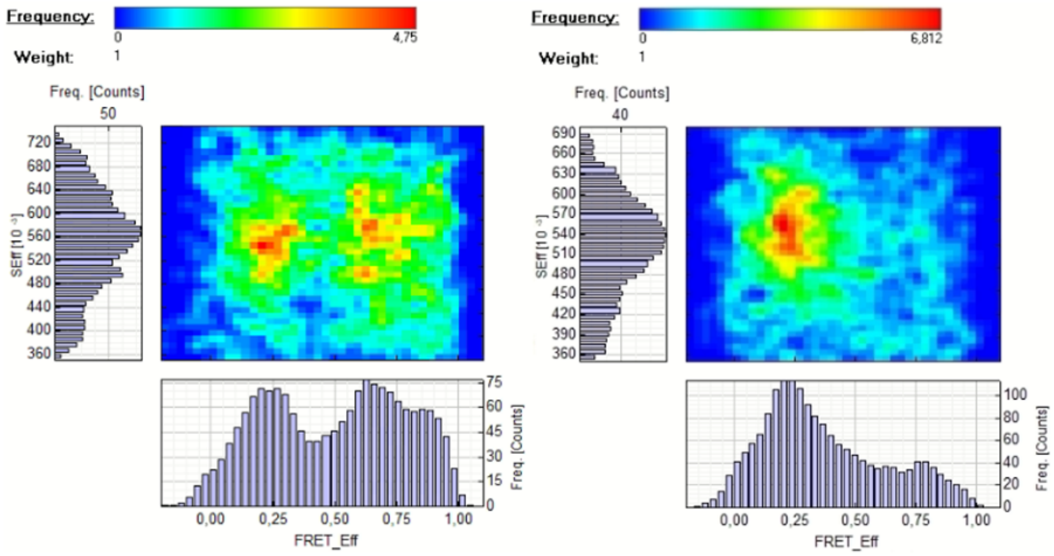


Supplementary Figure 11. FRET efficiency histograms of the asymmetric E33A-WT Hsp90 dimer (left) and the symmetric E33A-E33A dimer (right), both measured with 2 mM ATP. While the E33A-WT dimer exhibits a high-FRET efficiency, i.e., closed state, the E33A-E33A dimer is mainly found in the open state.

#### L. FRET-FCS data analysis

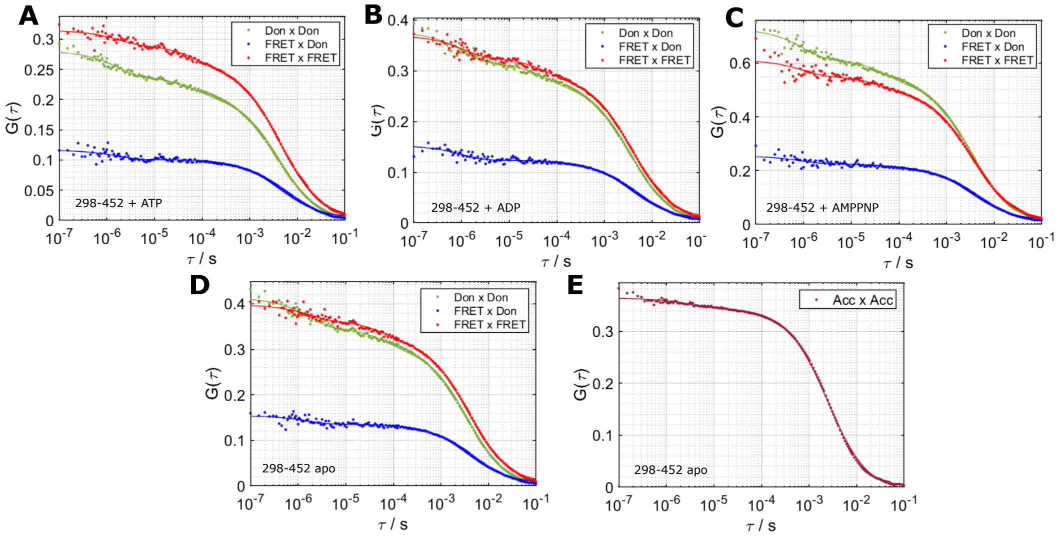


Supplementary Figure 12. FRET-FCS data and fits for FRET pair 298-452 of Hsp90. Autocorrelations DonxDon (green), FRETxFRET (red) and FRETxDon (blue) are shown for different nucleotide conditions: **a** ATP, **b** ADP, **c** AMPPNP and **d** apo. **e** Parallel and perpendicular parts of the acceptor signal after acceptor excitation are correlated for Hsp90 apo. The obtained AccxAcc correlation can be fit with model A, which includes simple 3D diffusion and a term for triplet kinetics only. The fact, that an additional kinetic term to fit AccxAcc is not necessary provides evidence, that we indeed see protein dynamics in a-d.

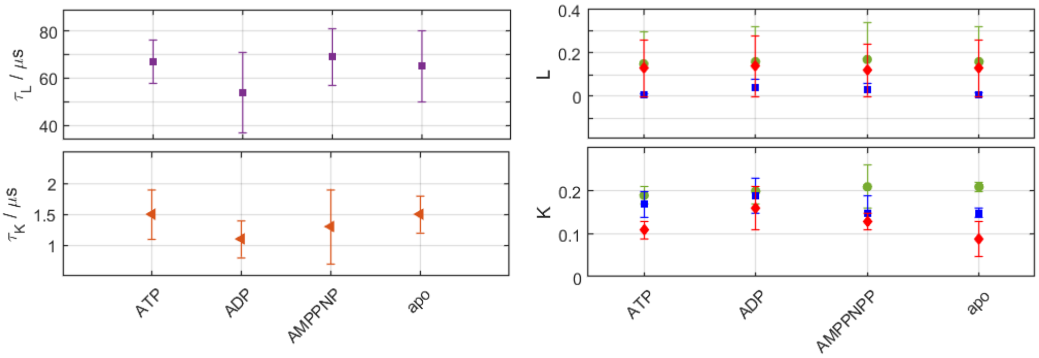


Supplementary Figure 13. FRET-FCS fit results for FRET pair 298-452 of Hsp90. Shown are the relaxation times for the fast (orange) and the slow kinetic mode (purple), $\tau_K$ and $\tau_L$, respectively and the corresponding weights of the kinetic modes $K$ and $L$ for different nucleotide conditions. For each weight three data points are shown which are the weights obtained from the DonxDon correlation (green cycle), from the FRETxFRET correlation (red diamond) and from the FRETxDon correlation (blue square). We could not observe a significant trend with respect to the nucleotide present. However, for the slow kinetic mode, the weights $L$ corresponding to FRETxDon correlation (blue squares) were always lower than those of DonxDon and FRETxFRET. This hints towards anti-correlated conformational dynamics at around 60 $\mu$s and correlated dynamics at around 1 $\mu$s.

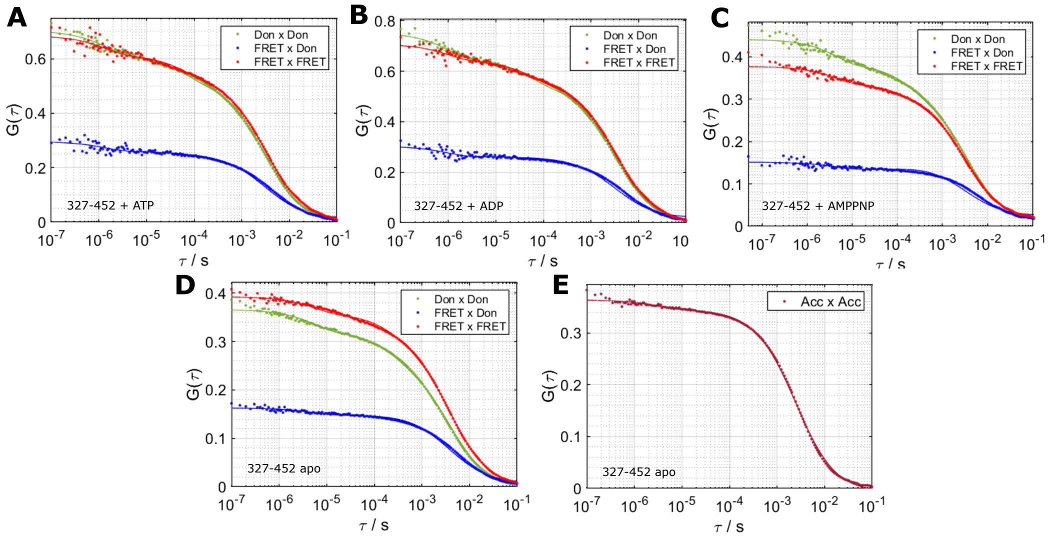


Supplementary Figure 14. FRET-FCS data and fits for FRET pair 327-452 of Hsp90. Autocorrelations DonxDon (green), FRETxFRET (red) and FRETxDon (blue) are shown for different nucleotide conditions: **a** ATP, **b** ADP, **c** AMPPNP and **d** apo. **e** Parallel and perpendicular parts of the acceptor signal after acceptor excitation are correlated for Hsp90 apo. The obtained AccxAcc correlation can be fit with model A, which includes simple 3D diffusion and a term for triplet kinetics only. The fact, that an additional kinetic term to fit AccxAcc is not necessary provides evidence, that we indeed see protein dynamics in a-d.

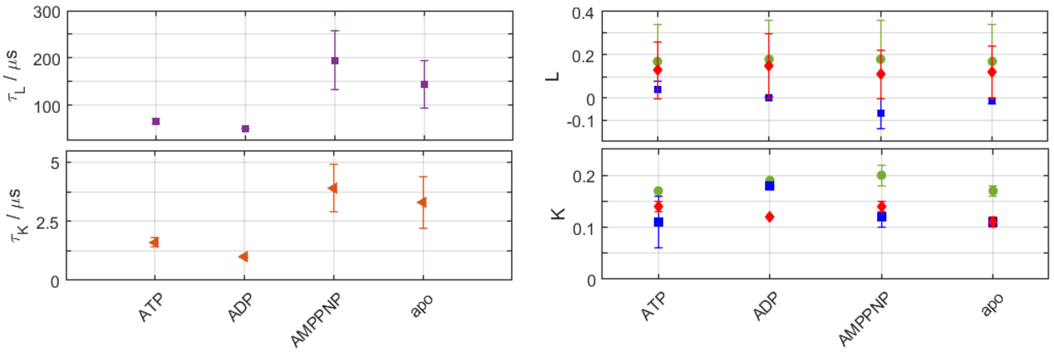


Supplementary Figure 15. FRET-FCS fit results for FRET pair 327-452 of Hsp90. Shown are the relaxation times for the fast (orange) and the slow kinetic mode (purple), $\tau_K$ and $\tau_L$, respectively and the corresponding weights of the kinetic modes $K$ and $L$ for different nucleotide conditions. For each weight three data points are shown which are the weights obtained from the DonxDon correlation (green cycle), from the FRETxFRET correlation (red diamond) and from the FRETxDon correlation (blue square). As for the FRET pair shown before, we could not observe a significant trend with respect to the nucleotide present. However, as for 298-452, the weights of the slow kinetic mode $L$ which correspond to the FRETxDon correlation (blue squares) were always lower than those of DonxDon and FRETxFRET. As well, this hints towards anti-correlated conformational dynamics which are here in the range of 50 to 200 $\mu$s and correlated dynamics at around 3 $\mu$s.

#### M. Targeted molecular dynamics simulations

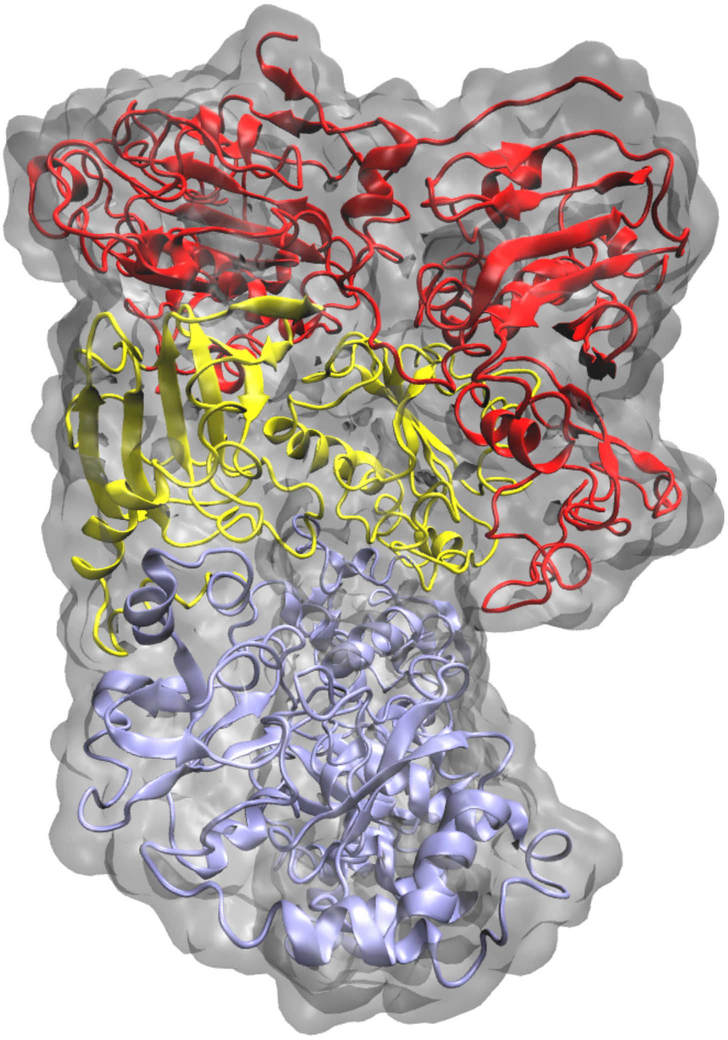

Supplementary Figure 16. Proposed model of closed state B based on nonequilibrium targeted MD simulations. Orientation and color code as in Fig. 1 in the main text. Protein surface in grey. The monomer on the right exhibits a kink at the M-C interface, causing the collapse of the central binding site and a rearrangement of N and M domains (see main text Fig. 8 for a sketch of the changes).

#### IV. SUPPLEMENTARY NOTES

##### A. Supplementary Note 1: AMPPNP parameters

[ moleculetype ]
;name nrexcl
AMPPNP 3

[ atoms ]
; nr type resi res atom cgnr charge mass ; qtot bondtype
1 O3 1 ANP O1G 1 -0.960834 16.00000 ; qtot -0.961
2 P 1 ANP PG 2 1.337801 30.97000 ; qtot 0.377
3 O3 1 ANP O2G 3 -0.960834 16.00000 ; qtot -0.584
4 O3 1 ANP O3G 4 -0.960834 16.00000 ; qtot -1.545
5 N 1 ANP N1B 5 -0.976401 14.01000 ; qtot -2.521
6 H 1 ANP H1B 6 0.370800 1.00800 ; qtot -2.150
7 P 1 ANP PB 7 1.644705 30.97000 ; qtot -0.506
8 O2 1 ANP O1B 8 -0.941001 16.00000 ; qtot -1.447
9 O2 1 ANP O2B 9 -0.941001 16.00000 ; qtot -2.388
10 OS 1 ANP O3A 10 -0.826801 16.00000 ; qtot -3.214
11 P 1 ANP PA 11 1.583802 30.97000 ; qtot -1.631
12 O2 1 ANP O1A 12 -0.906001 16.00000 ; qtot -2.537
13 O2 1 ANP O2A 13 -0.906001 16.00000 ; qtot -3.443
14 OS 1 ANP O5* 14 -0.580201 16.00000 ; qtot -4.023
15 CT 1 ANP C5* 15 0.198400 12.01000 ; qtot -3.824
16 H1 1 ANP H50 16 0.015700 1.00800 ; qtot -3.809
17 H1 1 ANP H51 17 0.015700 1.00800 ; qtot -3.793
18 CT 1 ANP C4* 18 0.095100 12.01000 ; qtot -3.698
19 H1 1 ANP H40 19 0.021700 1.00800 ; qtot -3.676
20 OS 1 ANP O4* 20 -0.446600 16.00000 ; qtot -4.123
21 CT 1 ANP C1* 21 0.248800 12.01000 ; qtot -3.874
22 H2 1 ANP H10 22 0.066700 1.00800 ; qtot -3.807

23 N* 1 ANP N9 23 -0.261000 14.01000 ; qtot -4.068

24 CK 1 ANP C8 24 0.543701 12.01000 ; qtot -3.525

25 H5 1 ANP H80 25 0.238800 1.00800 ; qtot -3.286

26 NB 1 ANP N7 26 -0.626101 14.01000 ; qtot -3.912

27 CB 1 ANP C5 27 -0.158200 12.01000 ; qtot -4.070

28 CA 1 ANP C6 28 0.626201 12.01000 ; qtot -3.444

29 N2 1 ANP N6 29 -0.909501 14.01000 ; qtot -4.353

30 H 1 ANP H60 30 0.381800 1.00800 ; qtot -3.972

31 H 1 ANP H61 31 0.381800 1.00800 ; qtot -3.590

32 NC 1 ANP N1 32 -0.804001 14.01000 ; qtot -4.394

33 CQ 1 ANP C2 33 0.591901 12.01000 ; qtot -3.802

34 H5 1 ANP H2 34 0.000100 1.00800 ; qtot -3.802

35 NC 1 ANP N3 35 -0.711001 14.01000 ; qtot -4.513

36 CB 1 ANP C4 36 0.414800 12.01000 ; qtot -4.098

37 CT 1 ANP C3* 37 0.098100 12.01000 ; qtot -4.000

38 H1 1 ANP H30 38 0.174700 1.00800 ; qtot -3.825

39 OH 1 ANP O3* 39 -0.674801 16.00000 ; qtot -4.500

40 HO 1 ANP H3* 40 0.496000 1.00800 ; qtot -4.004

41 CT 1 ANP C2* 41 0.083100 12.01000 ; qtot -3.921

42 H1 1 ANP H20 42 0.149700 1.00800 ; qtot -3.771

43 OH 1 ANP O2* 43 -0.653801 16.00000 ; qtot -4.425

44 HO 1 ANP H2* 44 0.425000 1.00800 ; qtot -4.000

[ bonds ]

; ai aj funct r k

1 2 1 ; O1G - PG

2 3 1 ; PG - O2G

2 4 1 ; PG - O3G

2 5 1 1.6696e-01 3.0711e+05 ; PG - N1B

5 6 1 1.0190e-01 3.2836e+05 ; N1B - H1B

5 7 1 1.6696e-01 3.0711e+05 ; N1B - PB

7 8 1 ; PB - O1B

7 9 1 ; PB - O2B

7 10 1 ; PB - O3A

10 11 1 ; O3A - PA

11 12 1 ; PA - O1A

11 13 1 ; PA - O2A

11 14 1 ; PA - O5*

14 15 1 ; O5* - C5*

15 16 1 ; C5* - H50

15 17 1 ; C5* - H51

15 18 1 ; C5* - C4*

18 19 1 ; C4* - H40

18 20 1 ; C4* - O4*

18 37 1 ; C4* - C3*

20 21 1 ; O4* - C1*

21 22 1 ; C1* - H10

21 23 1 ; C1* - N9

21 41 1 ; C1* - C2*

23 24 1 ; N9 - C8

23 36 1 ; N9 - C4

24 25 1 ; C8 - H80

24 26 1 ; C8 - N7

26 27 1 ; N7 - C5

27 28 1 ; C5 - C6

27 36 1 ; C5 - C4

28 29 1 ; C6 - N6

28 32 1 ; C6 - N1

29 30 1 ; N6 - H60

29 31 1 ; N6 - H61

32 33 1 ; N1 - C2

33 34 1 ; C2 - H2

33 35 1 ; C2 - N3

35 36 1 ; N3 - C4

37 38 1 ; C3* - H30

37 39 1 ; C3* - O3*

37 41 1 ; C3* - C2*

39 40 1 ; O3* - H3*

41 42 1 ; C2* - H20

41 43 1 ; C2* - O2*

43 44 1 ; O2* - H2*

[ pairs ]

; ai aj funct

1 6 1 ; O1G - H1B

1 7 1 ; O1G - PB

2 8 1 ; PG - O1B

2 9 1 ; PG - O2B

2 10 1 ; PG - O3A

3 6 1 ; O2G - H1B

3 7 1 ; O2G - PB

4 6 1 ; O3G - H1B

4 7 1 ; O3G - PB

5 11 1 ; N1B - PA

6 8 1 ; H1B - O1B

6 9 1 ; H1B - O2B

6 10 1 ; H1B - O3A

7 12 1 ; PB - O1A

7 13 1 ; PB - O2A

7 14 1 ; PB - O5*

8 11 1 ; O1B - PA

9 11 1 ; O2B - PA

10 15 1 ; O3A - C5*

11 16 1 ; PA - H50

11 17 1 ; PA - H51

11 18 1 ; PA - C4*

12 15 1 ; O1A - C5*

13 15 1 ; O2A - C5*

14 19 1 ; O5* - H40

14 20 1 ; O5* - O4*

14 37 1 ; O5* - C3*

15 21 1 ; C5* - C1*

15 38 1 ; C5* - H30

15 39 1 ; C5* - O3*

15 41 1 ; C5* - C2*

16 19 1 ; H50 - H40

16 20 1 ; H50 - O4*

16 37 1 ; H50 - C3*

17 19 1 ; H51 - H40

17 20 1 ; H51 - O4*

17 37 1 ; H51 - C3*

18 22 1 ; C4* - H10

18 23 1 ; C4* - N9

18 40 1 ; C4* - H3*

18 42 1 ; C4* - H20

18 43 1 ; C4* - O2*

19 21 1 ; H40 - C1*

19 38 1 ; H40 - H30

19 39 1 ; H40 - O3*

19 41 1 ; H40 - C2*

20 24 1 ; O4* - C8

20 36 1 ; O4* - C4

20 38 1 ; O4* - H30

20 39 1 ; O4* - O3*

20 42 1 ; O4* - H20

20 43 1 ; O4* - O2*

21 25 1 ; C1* - H80

21 26 1 ; C1* - N7

21 27 1 ; C1* - C5

21 35 1 ; C1* - N3

21 38 1 ; C1* - H30

21 39 1 ; C1* - O3*

21 44 1 ; C1* - H2*

22 24 1 ; H10 - C8

22 36 1 ; H10 - C4

22 37 1 ; H10 - C3*

22 42 1 ; H10 - H20

22 43 1 ; H10 - O2*

23 28 1 ; N9 - C6

23 33 1 ; N9 - C2

23 37 1 ; N9 - C3*

23 42 1 ; N9 - H20

23 43 1 ; N9 - O2*

24 28 1 ; C8 - C6

24 35 1 ; C8 - N3

24 41 1 ; C8 - C2*

25 27 1 ; H80 - C5

25 36 1 ; H80 - C4

26 29 1 ; N7 - N6

26 32 1 ; N7 - N1

26 35 1 ; N7 - N3

27 30 1 ; C5 - H60

27 31 1 ; C5 - H61

27 33 1 ; C5 - C2

28 34 1 ; C6 - H2

28 35 1 ; C6 - N3

29 33 1 ; N6 - C2

29 36 1 ; N6 - C4

30 32 1 ; H60 - N1

31 32 1 ; H61 - N1

32 36 1 ; N1 - C4

34 36 1 ; H2 - C4

36 41 1 ; C4 - C2*

37 44 1 ; C3* - H2*

38 40 1 ; H30 - H3*

38 42 1 ; H30 - H20

38 43 1 ; H30 - O2*

39 42 1 ; O3* - H20

39 43 1 ; O3* - O2*

40 41 1 ; H3* - C2*

42 44 1 ; H20 - H2*

[ angles ]

; ai aj ak funct theta cth

1 2 3 1 ; O1G - PG - O2G

1 2 4 1 ; O1G - PG - O3G

1 2 5 1 ; O1G - PG - N1B

2 5 6 1 1.1432e+02 4.6568e+02 ; PG - N1B - H1B

2 5 7 1 ; PG - N1B - PB

3 2 4 1 ; O2G - PG - O3G

3 2 5 1 ; O2G - PG - N1B

4 2 5 1 ; O3G - PG - N1B

5 7 8 1 ; N1B - PB - O1B

5 7 9 1 ; N1B - PB - O2B

5 7 10 1 ; N1B - PB - O3A

6 5 7 1 1.1432e+02 4.6568e+02 ; H1B - N1B - PB

7 10 11 1 ; PB - O3A - PA

8 7 9 1 ; O1B - PB - O2B

8 7 10 1 ; O1B - PB - O3A

9 7 10 1 ; O2B - PB - O3A

10 11 12 1 ; O3A - PA - O1A

10 11 13 1 ; O3A - PA - O2A

10 11 14 1 ; O3A - PA - O5*

11 14 15 1 ; PA - O5* - C5*

12 11 13 1 ; O1A - PA - O2A

12 11 14 1 ; O1A - PA - O5*

13 11 14 1 ; O2A - PA - O5*

14 15 16 1 ; O5* - C5* - H50

14 15 17 1 ; O5* - C5* - H51

14 15 18 1 ; O5* - C5* - C4*

15 18 19 1 ; C5* - C4* - H40

15 18 20 1 ; C5* - C4* - O4*

15 18 37 1 ; C5* - C4* - C3*

16 15 17 1 ; H50 - C5* - H51

16 15 18 1 ; H50 - C5* - C4*

17 15 18 1 ; H51 - C5* - C4*

18 20 21 1 ; C4* - O4* - C1*

18 37 38 1 ; C4* - C3* - H30

18 37 39 1 ; C4* - C3* - O3*

18 37 41 1 ; C4* - C3* - C2*

19 18 20 1 ; H40 - C4* - O4*

19 18 37 1 ; H40 - C4* - C3*

20 18 37 1 ; O4* - C4* - C3*

20 21 22 1 ; O4* - C1* - H10

20 21 23 1 ; O4* - C1* - N9

20 21 41 1 ; O4* - C1* - C2*

21 23 24 1 ; C1* - N9 - C8

21 23 36 1 ; C1* - N9 - C4

21 41 37 1 ; C1* - C2* - C3*

21 41 42 1 ; C1* - C2* - H20

21 41 43 1 ; C1* - C2* - O2*

22 21 23 1 ; H10 - C1* - N9

22 21 41 1 ; H10 - C1* - C2*

23 21 41 1 ; N9 - C1* - C2*

23 24 25 1 ; N9 - C8 - H80

23 24 26 1 ; N9 - C8 - N7

23 36 27 1 ; N9 - C4 - C5

23 36 35 1 ; N9 - C4 - N3

24 23 36 1 ; C8 - N9 - C4

24 26 27 1 ; C8 - N7 - C5

25 24 26 1 ; H80 - C8 - N7

26 27 28 1 ; N7 - C5 - C6

26 27 36 1 ; N7 - C5 - C4

27 28 29 1 ; C5 - C6 - N6

27 28 32 1 ; C5 - C6 - N1

27 36 35 1 ; C5 - C4 - N3

28 27 36 1 ; C6 - C5 - C4

28 29 30 1 ; C6 - N6 - H60

28 29 31 1 ; C6 - N6 - H61

28 32 33 1 ; C6 - N1 - C2

29 28 32 1 ; N6 - C6 - N1

30 29 31 1 ; H60 - N6 - H61

32 33 34 1 ; N1 - C2 - H2

32 33 35 1 ; N1 - C2 - N3

33 35 36 1 ; C2 - N3 - C4

34 33 35 1 ; H2 - C2 - N3

37 39 40 1 ; C3* - O3* - H3*

37 41 42 1 ; C3* - C2* - H20

37 41 43 1 ; C3* - C2* - O2*

38 37 39 1 ; H30 - C3* - O3*

38 37 41 1 ; H30 - C3* - C2*

39 37 41 1 ; O3* - C3* - C2*

41 43 44 1 ; C2* - O2* - H2*

42 41 43 1 ; H20 - C2* - O2*

[ dihedrals ] ; propers

; treated as RBs in GROMACS to use combine multiple AMBER torsions per quartet

; i j k l func C0 C1 C2 C3 C4 C5

1 2 5 6 3 25.10400 0.00000 -25.10400 0.00000 0.00000 0.00000 ; O1G- PG- N1B- H1B

1 2 5 7 9 ; O1G- PG- N1B- PB

2 5 7 8 9 ; PG- N1B- PB- O1B

2 5 7 9 9 ; PG- N1B- PB- O2B

2 5 7 10 9 ; PG- N1B- PB- O3A

3 2 5 6 3 25.10400 0.00000 -25.10400 0.00000 0.00000 0.00000 ; O2G- PG- N1B- H1B

3 2 5 7 9 ; O2G- PG- N1B- PB

4 2 5 6 3 25.10400 0.00000 -25.10400 0.00000 0.00000 0.00000 ; O3G- PG- N1B- H1B

4 2 5 7 9 ; O3G- PG- N1B- PB

5 7 10 11 9 ; N1B- PB- O3A- PA

6 5 7 8 3 25.10400 0.00000 -25.10400 0.00000 0.00000 0.00000 ; H1B- N1B- PB- O1B

6 5 7 9 3 25.10400 0.00000 -25.10400 0.00000 0.00000 0.00000 ; H1B- N1B- PB- O2B

6 5 7 10 3 25.10400 0.00000 -25.10400 0.00000 0.00000 0.00000 ; H1B- N1B- PB- O3A

7 10 11 12 9 ; PB- O3A- PA- O1A

7 10 11 13 9 ; PB- O3A- PA- O2A

7 10 11 14 9 ; PB- O3A- PA- O5*

8 7 10 11 9 ; O1B- PB- O3A- PA

9 7 10 11 9 ; O2B- PB- O3A- PA

10 11 14 15 9 ; O3A- PA- O5*- C5*

11 14 15 16 9 ; PA- O5*- C5*- H50

11 14 15 17 9 ; PA- O5*- C5*- H51

11 14 15 18 9 ; PA- O5*- C5*- C4*

12 11 14 15 9 ; O1A- PA- O5*- C5*

13 11 14 15 9 ; O2A- PA- O5*- C5*

14 15 18 19 9 ; O5*- C5*- C4*- H40

14 15 18 20 9 ; O5*- C5*- C4*- O4*

14 15 18 37 9 ; O5*- C5*- C4*- C3*

15 18 20 21 9 ; C5*- C4*- O4*- C1*

15 18 37 38 9 ; C5*- C4*- C3*- H30

15 18 37 39 9 ; C5*- C4*- C3*- O3*

15 18 37 41 9 ; C5*- C4*- C3*- C2*

16 15 18 19 9 ; H50- C5*- C4*- H40

16 15 18 20 9 ; H50- C5*- C4*- O4*

16 15 18 37 9 ; H50- C5*- C4*- C3*

17 15 18 19 9 ; H51- C5*- C4*- H40

17 15 18 20 9 ; H51- C5*- C4*- O4*

17 15 18 37 9 ; H51- C5*- C4*- C3*

18 20 21 22 9 ; C4*- O4*- C1*- H10

18 20 21 23 9 ; C4*- O4*- C1*- N9

18 20 21 41 9 ; C4*- O4*- C1*- C2*

18 37 39 40 9 ; C4*- C3*- O3*- H3*

18 37 41 21 9 ; C4*- C3*- C2*- C1*

18 37 41 42 9 ; C4*- C3*- C2*- H20

18 37 41 43 9 ; C4*- C3*- C2*- O2*

19 18 20 21 9 ; H40- C4*- O4*- C1*

19 18 37 38 9 ; H40- C4*- C3*- H30

19 18 37 39 9 ; H40- C4*- C3*- O3*

19 18 37 41 9 ; H40- C4*- C3*- C2*

20 18 37 38 9 ; O4*- C4*- C3*- H30

20 18 37 39 9 ; O4*- C4*- C3*- O3*

20 18 37 41 9 ; O4*- C4*- C3*- C2*

20 21 23 24 9 ; O4*- C1*- N9- C8

20 21 23 36 9 ; O4*- C1*- N9- C4

20 21 41 37 9 ; O4*- C1*- C2*- C3*

20 21 41 42 9 ; O4*- C1*- C2*- H20

20 21 41 43 9 ; O4*- C1*- C2*- O2*

21 20 18 37 9 ; C1*- O4*- C4*- C3*

21 23 24 25 9 ; C1*- N9- C8- H80

21 23 24 26 9 ; C1*- N9- C8- N7

21 23 36 27 9 ; C1*- N9- C4- C5

21 23 36 35 9 ; C1*- N9- C4- N3

21 41 37 38 9 ; C1*- C2*- C3*- H30

21 41 37 39 9 ; C1*- C2*- C3*- O3*

21 41 43 44 9 ; C1*- C2*- O2*- H2*

22 21 23 24 9 ; H10- C1*- N9- C8

22 21 23 36 9 ; H10- C1*- N9- C4

22 21 41 37 9 ; H10- C1*- C2*- C3*

22 21 41 42 9 ; H10- C1*- C2*- H20

22 21 41 43 9 ; H10- C1*- C2*- O2*

23 21 41 37 9 ; N9- C1*- C2*- C3*

23 21 41 42 9 ; N9- C1*- C2*- H20

23 21 41 43 9 ; N9- C1*- C2*- O2*

23 24 26 27 9 ; N9- C8- N7- C5

23 36 27 26 9 ; N9- C4- C5- N7

23 36 27 28 9 ; N9- C4- C5- C6

23 36 35 33 9 ; N9- C4- N3- C2

24 23 21 41 9 ; C8- N9- C1*- C2*

24 23 36 27 9 ; C8- N9- C4- C5

24 23 36 35 9 ; C8- N9- C4- N3

24 26 27 28 9 ; C8- N7- C5- C6

24 26 27 36 9 ; C8- N7- C5- C4

25 24 23 36 9 ; H80- C8- N9- C4

25 24 26 27 9 ; H80- C8- N7- C5

26 24 23 36 9 ; N7- C8- N9- C4

26 27 28 29 9 ; N7- C5- C6- N6

26 27 28 32 9 ; N7- C5- C6- N1

26 27 36 35 9 ; N7- C5- C4- N3

27 28 29 30 9 ; C5- C6- N6- H60

27 28 29 31 9 ; C5- C6- N6- H61

27 28 32 33 9 ; C5- C6- N1- C2

27 36 35 33 9 ; C5- C4- N3- C2

28 27 36 35 9 ; C6- C5- C4- N3

28 32 33 34 9 ; C6- N1- C2- H2

28 32 33 35 9 ; C6- N1- C2- N3

29 28 27 36 9 ; N6- C6- C5- C4

29 28 32 33 9 ; N6- C6- N1- C2

30 29 28 32 9 ; H60- N6- C6- N1

31 29 28 32 9 ; H61- N6- C6- N1

32 28 27 36 9 ; N1- C6- C5- C4

32 33 35 36 9 ; N1- C2- N3- C4

34 33 35 36 9 ; H2- C2- N3- C4

36 23 21 41 9 ; C4- N9- C1*- C2*

37 41 43 44 9 ; C3*- C2*- O2*- H2*

38 37 39 40 9 ; H30- C3*- O3*- H3*

38 37 41 42 9 ; H30- C3*- C2*- H20

38 37 41 43 9 ; H30- C3*- C2*- O2*

39 37 41 42 9 ; O3*- C3*- C2*- H20

39 37 41 43 9 ; O3*- C3*- C2*- O2*

40 39 37 41 9 ; H3*- O3*- C3*- C2*

42 41 43 44 9 ; H20- C2*- O2*- H2*

[ dihedrals ] ; impropers

; ai aj ak al funct c0 c1 c2 c3

2 5 7 6 4 180.00 4.60240 1

21 23 36 24 4 180.00 4.60240 1

25 24 23 26 4 180.00 4.60240 1

27 32 28 29 4 180.00 4.60240 1

28 30 29 31 4 180.00 4.60240 1

32 35 33 34 4 180.00 4.60240 1

#### B. Supplementary Note 2: $H_2PO_4^-$ parameters

[ moleculetype ]

;name nrexcl

PO4 3

[ atoms ]

; nr type resi res atom cgnr charge mass ; qtot bondtype

1 P 1 PO4 P 1 1.427802 30.97000 ; qtot 1.428

2 O3 1 PO4 O1 2 -0.849501 16.00000 ; qtot 0.578

3 O3 1 PO4 O2 3 -0.849501 16.00000 ; qtot -0.271

4 OH 1 PO4 O3 4 -0.784401 16.00000 ; qtot -1.056

5 HO 1 PO4 H03 5 0.420000 1.00800 ; qtot -0.636

6 OH 1 PO4 O4 6 -0.784401 16.00000 ; qtot -1.420

7 HO 1 PO4 H04 7 0.420000 1.00800 ; qtot -1.000

[ bonds ]

; ai aj funct r k

1 2 1 1.4866e-01 4.0125e+05 ; P - O1

1 3 1 1.4866e-01 4.0125e+05 ; P - O2

1 4 1 1.6149e-01 2.7648e+05 ; P - O3

1 6 1 1.6149e-01 2.7648e+05 ; P - O4

4 5 1 9.7300e-02 3.1079e+05 ; O3 - H03

6 7 1 9.7300e-02 3.1079e+05 ; O4 - H04

[ exclusions ]

; ai aj funct

2 5 1 ; O1 - H03

2 7 1 ; O1 - H04

3 5 1 ; O2 - H03

3 7 1 ; O2 - H04

4 7 1 ; O3 - H04

6 5 1 ; O4 - H03

[ angles ]

; ai aj ak funct theta cth

1 4 5 1 1.1008e+02 4.7087e+02 ; P - O3 - H03

1 6 7 1 1.1008e+02 4.7087e+02 ; P - O4 - H04

2 1 3 1 1.1580e+02 3.8351e+02 ; O1 - P - O2

2 1 4 1 1.1521e+02 3.6736e+02 ; O1 - P - O3

2 1 6 1 1.1521e+02 3.6736e+02 ; O1 - P - O4

3 1 4 1 1.1521e+02 3.6736e+02 ; O2 - P - O3

3 1 6 1 1.1521e+02 3.6736e+02 ; O2 - P - O4

4 1 6 1 1.0269e+02 3.7489e+02 ; O3 - P - O4

[ dihedrals ]

;improper to keep the phosphate tetrahedron stable in initial relaxation run

; 2 1 3 4 2 125 1000 ; parameters based on JCC,7,(1986),230

; 2 1 3 6 2 -125 1000 ; parameters based on JCC,7,(1986),230

; 2 1 4 6 2 118 1000 ; parameters based on JCC,7,(1986),230

; 3 1 4 6 2 -118 1000 ; parameters based on JCC,7,(1986),230

; propers

; treated as RBs in GROMACS to use combine multiple AMBER torsions per quartet

; i j k l func C0 C1 C2 C3 C4 C5

2 1 4 5 3 2.23147 6.69440 0.00000 -8.92587 0.00000 0.00000 ; O1- P- O3- H03

2 1 6 7 3 2.23147 6.69440 0.00000 -8.92587 0.00000 0.00000 ; O1- P- O4- H04

3 1 4 5 3 2.23147 6.69440 0.00000 -8.92587 0.00000 0.00000 ; O2- P- O3- H03

3 1 6 7 3 2.23147 6.69440 0.00000 -8.92587 0.00000 0.00000 ; O2- P- O4- H04

4 1 6 7 3 2.23147 6.69440 0.00000 -8.92587 0.00000 0.00000 ; O3- P- O4- H04

6 1 4 5 3 2.23147 6.69440 0.00000 -8.92587 0.00000 0.00000 ; O4- P- O3- H03

#### C. Supplementary Note 3: Additional simulation parameters

new angles:

O3 P OH 1 108.230 376.560

O2 P N 1 108.230 836.800

O3 P N 1 108.230 836.800

OH P OH 1 102.600 376.560

N P OS 1 102.600 376.560

new ADP/ATP dihedrals:

H1 CT OS P 9 0.0 0.4389 3.000 ; J Comp Chem 2003, 24, 1016-1025

O2 P OS CT 9 0.0 4.9282 -3.000 ; J Comp Chem 2003, 24, 1016-1025

O2 P OS CT 9 0.0 -3.3942 2.000 ; J Comp Chem 2003, 24, 1016-1025

CT OS P OS 9 0.0 -19.0608 1.0 ; J Comp Chem 2003, 24, 1016-1025

O2 P OS P 9 0.0 -2.9636 2.0 ; J Comp Chem 2003, 24, 1016-1025

O2 P N P 9 0.0 -2.9636 2.0 ; copied from above

O3 P OS P 9 0.0 -1.0659 3.0 ; J Comp Chem 2003, 24, 1016-1025

O3 P N P 9 0.0 -1.0659 3.0 ; copied from above

P OS P OS 9 0.0 3.7495 1.0 ; J Comp Chem 2003, 24, 1016-1025

P N P OS 9 0.0 3.7495 1.0 ; copied from above
